# Diagnostic Evidence Gauge of Spatial Transcriptomics (DEGAS): Using transfer learning to map clinical data to spatial transcriptomics in prostate cancer

**DOI:** 10.1101/2023.04.21.537852

**Authors:** Justin L. Couetil, Ziyu Liu, Ahmed K. Alomari, Jie Zhang, Kun Huang, Travis S. Johnson

## Abstract

**Background:** Spatial and single-cell transcriptomics have revealed significant heterogeneity in tumor and normal tissues. Each approach has its advantages: The Visium platform for spatial transcriptomics (ST) offers lower resolution than single-cell analysis, but histology enables the examination of cell morphology, tissue architecture, and potential cell-cell interactions. Single-cell transcriptomics (SC) provides high resolution, but manual cell-type annotation depends on incomplete scientific knowledge from heterogeneous experiments. When investigating poorly defined phenomena, such as the transition from normal tissue to cancer and metaplasia, researchers might overlook critical and unexpected findings in downstream analysis if they rely on pre-existing annotations to determine cell types, particularly in the context of phenotypic plasticity.

**Results:** We employ our deep-transfer learning framework, DEGAS, to identify benign morphology glands in normal prostate tissue that are associated with poor progression-free survival in cancer patients and exhibit transcriptional signatures of carcinogenesis and de-differentiation. We confirm this finding in an additional ST dataset and use novel published methods to integrate SC data, showing that cells annotated as cancerous in the SC data map to regions of benign glands in another dataset. We pinpoint several genes, primarily MSMB, with expression closely correlated with progression-free survival scores, which are known markers of de-differentiation, and attribute their expression specifically to luminal epithelia, which are the presumed origin of most prostatic cancers.

**Discussion:** Our work shows that morphologically normal epithelia can have transcriptional signatures like that of frank cancer, and that these tissues are associated with poor progression-free survival. We also highlight a critical gap in single-cell workflows: annotating continuous transitional phenomena like carcinogenesis with discrete labels can result in incomplete conclusions. Two approaches can help mitigate this issue: Tools like DEGAS and Scissor can provide a disease-association score for SC and ST data, independent of cell type and histology. Additionally, researchers should adopt a bidirectional approach, transferring histological labels from ST data to SC data using tools like RCTD, rather than only using SC cell-type assignments to annotate ST data. Employed together, these methods can offer valuable histology and disease-related information to better define tissue subtypes, especially epithelial cells in the process of carcinogenesis.

**Conclusions:** DEGAS is a vital tool for generating clinically-oriented hypotheses from SC and ST data, which are heterogeneous, information-rich assays. In this study, we identify potential signatures of carcinogenesis in morphologically benign epithelia, which may be the precursors to cancer and high-grade pre-malignant lesions. Validating these genes as a panel may help identify patients at high risk for future cancer development, recurrence, and assist researchers in studying the biology of early carcinogenesis by detecting metaplastic changes before they are morphologically identifiable.

## INTRODUCTION

Prostate cancer is the most common cancer specific to the male sex, and second most fatal cancer among men in most regions of the world (1). Treatment involves a combination of radiotherapy, surgery, and androgen deprivation therapy (2). There is also a frequent recurrence after surgery: In a retrospective cohort of 450 men with median follow up time of 8 years, ∼29.8% of patients developed metastasis after radical prostatectomy (3). Prostate cancer prevention and disease monitoring currently relies mainly on tracking the level of prostate specific antigen (PSA*/KLK-3*). This is a protease secreted by glandular tissues of the prostate and therefore generally rises in the development of progression of carcinoma (4). PSA screening, however, has not consistently been associated with an improvement in cancer-specific mortality, and in-fact, is associated with >50% overdiagnosis and common overtreatment (5).

Many cancers have known morphological precursors. This is the basis of cancer screening. Pre-malignant lesions may be grossly visible to the naked eye (i.e., a suspicious polyp during colonoscopy). Some lesions demand careful microscopic inspection and immunohistochemistry, like melanoma arising in the background of a benign pigmented nevus (6). The process of carcinogenesis involves metaplasia, a physiologic process whereby cells de-differentiate and lose polarity to migrate and heal injured tissue. In the context of cancer, metaplasia is viewed as a precursor to dysplasia (7). Dysplasia is the atypical growth of cells. Dysplasia is a precursor to *intraepithelial neoplasia*, which is in turn thought as a precursor to *carcinoma in-situ.* The latter is malignant neoplasia (new growth) contained by the basement membrane. The final diagnosis of invasive carcinoma is defined by the migration of neoplastic cells beyond the basement membrane and into nearby tissues. Cancer can hijack components of the metaplastic program for migration and metastasis (8).

Prostatic Intraepithelial Neoplasia (PIN), a precursor to prostate carcinoma, is defined by nuclear atypia and cell proliferations within ducts/glands that are contained within the basement membrane. PIN is a morphological diagnosis and associated with genomic aberrations that are shared between adjacent tumor (9). PIN increases in grade, size, and prevalence throughout life (10). In the United States, the prevalence of PIN within prostate biopsies is between 4% and 16% (11). Given that transcription will proceed protein expression, thus morphological changes, we hypothesize that there are transcriptional precursors to morphological changes of PIN and metaplasia, a part of the epithelial-to-mesenchyme transition. The epithelial-to-mesenchyme transition (EMT) is an incompletely understood continuum of genetic and epigenetic changes whereby cells are pushed by internal and external signals to lose their differentiated structure, function, and polarity. This favors a mesenchymal state better suited to tissue repair (12). The EMT program can be hijacked in the initiation, progression, metastasis, and chemotherapy resistance of cancers. Markers for metaplastic tissue are therefore of direct interest to aid cancer screening, basic science, and therapeutic development (13).

The gradual transitional nature of the EMT makes it extremely difficult to define distinct cell types, so attributing simple “normal” or “cancer” labels to transitional cells is fraught with potential for mislabeling. This is especially important for SC transcriptomics, where cell morphology and location in the tissue is not observable. Well-documented “field effects” provided evidence for space-varying pro-inflammatory environments in adjacent-normal tissue (14) and benign morphology cells can have shared epigenetic changes with adjacent tumors, demonstrating common lineage (15). This has important implications for clinical care: Firstly, “recurrence” of tumors which were surgically removed may not only be due to residual tumor (i.e., margin status), but also be related to benign morphology tissue left behind in tissue-sparing surgery that is predisposed to becoming cancer. This has been observed in breast cancer, where some local recurrences are similar to the original tumor and some seem to be distinct primaries (16). Secondly, existing screening tools in prostate cancer are poorly predictive (5), so understanding early carcinogenesis may help clinicians and researchers identify patients at high risk for developing cancer.

Herein, we report that DEGAS, a deep-transfer learning model, highlights specific regions of morphologically benign glandular prostatic tissues associated with poor progression-free survival in prostate cancer patients. These regions show transcriptional signatures associated with inflammation, metaplasia, and cancer, and colocalize with high grade prostate cancer tissue in clustering experiments. Risk scores are tightly correlated with overall transcriptional activity, but the expression of several epithelial marker genes is specifically lost in the highest-risk most transcriptionally active regions. Though trained on TCGA data, the DEGAS model predictions provide nuance that is not apparent in the TCGA transcriptomic and Human Protein Atlas proteomic datasets. We then analyzed a separate dataset, using RTCD and C-SIDE tools to integrate an annotated single-cell dataset, deconvoluting cell identities at each spot in the spatial transcriptomic arrays, and identifying cell-type specific differential gene expression that is associated with poor progression-free survival (**Fig. 1**). This analysis revealed that SC clusters annotated as tumor tissue localize to histologically benign regions of the ST data. Taken together, our results demonstrate that many “normal” epithelial cells are not truly normal, may be associated with cancer progression, and that using discrete cell-type annotations to define cells in transition provides an incomplete picture of the tumor and microenvironment. DEGAS is therefore a useful tool to identify disease-associated tissue and to define and refine SC clusters. We are using these results to develop gene panels that triage patients at risk for developing prostate cancer and recurrence.

**Figure 1:**
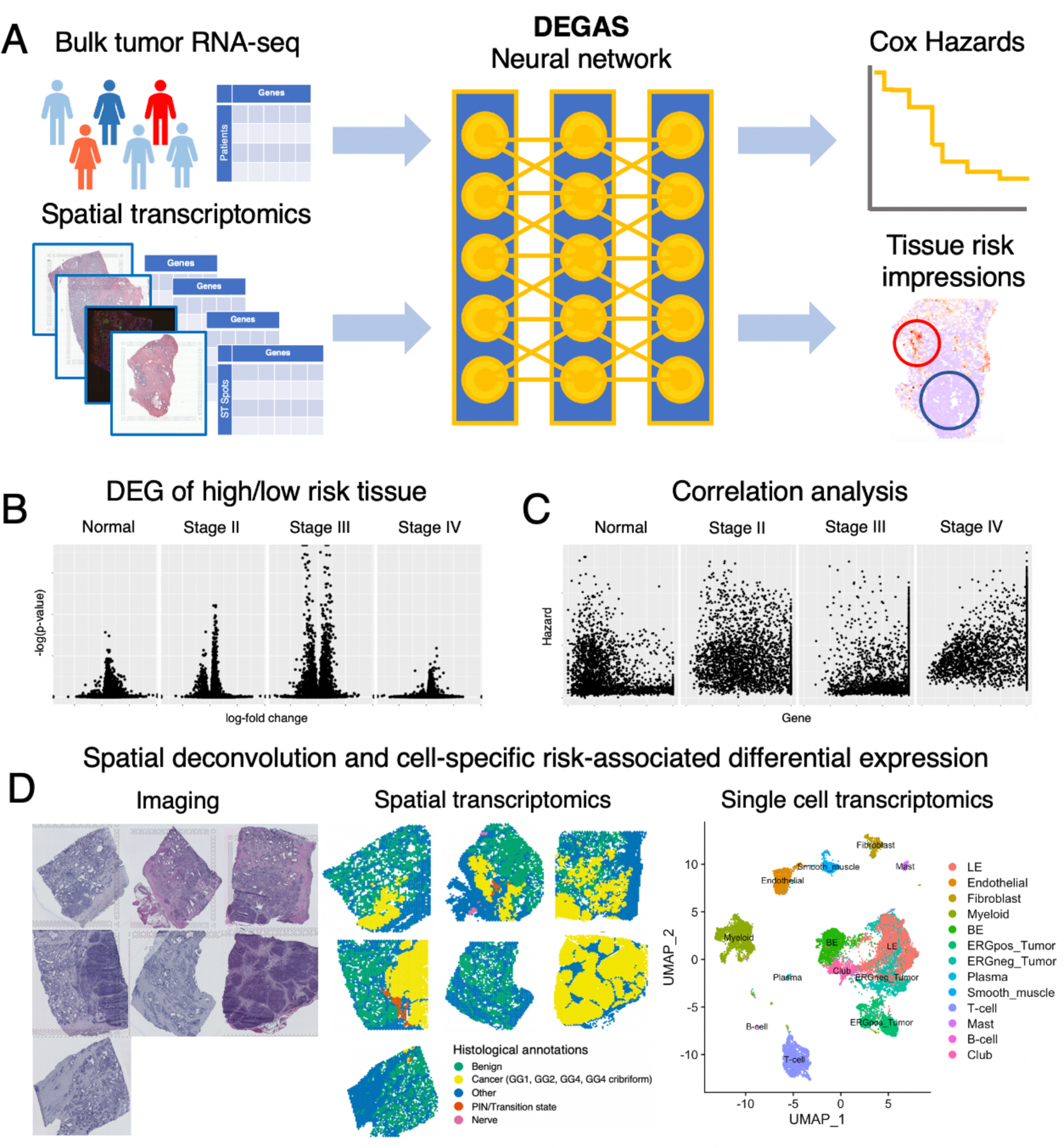
DEGAS workflow. **DEGAS workflow: A.** Patient bulk tumor RNA-seq is used to predict clinical outcomes and phenotypic information using a deep-learning framework that simultaneously matches ST gene expression distribution to that of patients. By aligning gene expression distributions across these two datatypes, DEGAS can then assign (“transfer”) risk scores to the spatial transcriptomic arrays, highlighting high-risk tissue. **B.** RNA expression at high and low-risk tissue are compared and **(C)** consistent correlations between gene expression and risk are identified. **D.** Morphologically benign glands with malignant transcriptional profiles are identified in an additional dataset. Integration with single cell data allows for identification of cell-type specific expression changes associated with poor progression-free survival, and that SC clusters annotated as cancer localize to histologically benign regions of ST data.

## METHODS

### Spatial Transcriptomic & RNA-seq data processing and analysis

The 10x Genomics data repository provided all spatial transcriptomic samples used in this work (17, 18). We use a Breast Ductal Carcinoma in-Situ (DCIS) sample, and four prostate tissues: Normal tissue, and Stage II, Stage III, and Stage IV adenocarcinoma. Patient-level RNA sequencing and clinical data for prostate cancer is derived from the TCGA and downloaded from *FireBrowse* (19). Gene TPMs for normal prostate tissue were downloaded from GTEX analysis v8 (20). We used this breast cancer dataset because it had histological labels, and the histology is easily visible at a low resolution.

### DEGAS framework

DEGAS is a neural network built with python TensorFlow (21) and wrapped with a convenient R interface. The framework is fully elaborated in the original publication (22). Briefly, DEGAS uses the maximum mean discrepancy loss to align the distributions of two different datasets in latent space. Once these datasets are aligned, class labels and patient outcomes (e.g., mutational status and survival) from the patient RNA-seq “training set” can be mapped on to similar observations in the spatial transcriptomic “test set”. DEGAS can be downloaded from GitHub at https://github.com/tsteelejohnson91/DEGAS.

### DEGAS pre-processing, feature selection, and exploratory datasets

All RNA data was processed for analysis in DEGAS with the built-in function *preprocessCounts*. This applies a log-2 transform, Z-score normalization, and scales RNA counts from zero to one for each gene. Predictions were all performed with a DEGAS DenseNet configuration with three hidden layers and five-fold bootstrap aggregation (default settings). The predicted hazard outputs for ST data were scaled by the median. This provides negative and positive values. “High-risk” is defined as a hazard greater than zero and low-risk is a hazard less than zero. The prostate DEGAS model was trained on a concatenated matrix of all four ST samples; therefore, the risk impression from the model can be compared across prostate cancer samples directly. For feature selection, the *var* function in base R version 4.0.4 (23) is used to find the 200 most variable genes among the patient-level RNA-seq datasets that overlap with each ST dataset (Supp. Figure 1). These genes were used as features in DEGAS BlankCox models in the prostate cancer analysis. Pooling results from sequentially larger and larger features sets (i.e., 10, 20, 40, 80, …, 200 genes) allows for regularization of the DEGAS hazard predictions (Supp. Figure 2). The “Blank” denomination refers to there being no class labels provided for the spatial transcriptomic samples. For the baseline comparison demonstrating resistance to data sparsity, we used a ClassCox and BlankCox model for illustrative purposes. Feature selection for this baseline comparison is further discussed in the relevant section below.

For the exploratory analysis in prostate cancer manuscript, four 10x genomics prostate cancer ST arrays were used (18). For external validation, seven prostate spatial transcriptomics samples were gathered from Erickson et al (15). These samples are different sections of the prostate from a single patient. These seven regions are histologically heterogenous, and the original authors identified morphologically benign glands with copy-number changes, showed shared tumor-normal tissue phylogenies, and thus provide compelling evidence that CNV’s are an early step in carcinogenesis. The dataset is thus appropriate for our analysis. All ST data followed the previously-described normalization procedures.

### Differential gene expression & correlation of DEGAS-Hazard with gene expression

Li, Ge (24), demonstrated that the Wilcoxon test can be used for differential expression to avoid false positives, while not unduly decreasing true positives. The log2 fold-change in mean expression between comparison groups is calculated to generate volcano plots. P-values are adjusted for false discovery with the Benjamini-Hochberg method available in the R package *stats v4.1.1 (25)*. The Spearman correlation between DEGAS risk scores and gene expression were calculated for every ST prostate sample, using the *cor* function from the same R *stats* package.

### Functional enrichment

We used the 100 most significantly differentially expressed genes with FDR-adjusted B&H p-values smaller than 1E-3, whose log2 fold-change was not infinite, for functional enrichment in Toppgene (26). Tissue regions being compared for differential expression are explained in the relevant experiment. Reported P-values from Toppgene are FDR B&H adjusted. There is often significant overlap in the meaning of GO Enrichment Terms. In these cases, one of a few similar ontologies were included in this manuscript for clarity.

### Dimensionality reduction with UMAP and t-SNE

R packages *umap* v0.2.9.0 (27) and *Rtsne* v0.15 (28) with default settings were used to cluster ST spots based on the 3000 most variable genes whose expression was normalized using the *preprocessCounts* function described earlier. This normalization procedure was carried out for each of the four samples individually, and not across all four samples.

### RNA-seq analysis

DEGAS is trained on TCGA data, but we found that it detects nuance in ST and SC data that is not easily observable in the TCGA data itself. To demonstrate this, we compared tumor and normal RNA-seq data from the TCGA and GTEX respectively (20). Directly comparing transcript counts from RNA-seq of disparate datasets that were normalized with different technique is technically very challenging and prone to bias (29). Instead, we instead focus on analyzing the statistics of the log10 transformed gene expression distribution: if a gene is among the most highly expressed genes in a cancer dataset, but among the lowest expressed genes in the normal dataset (and visa-versa), this swing in expression may help us identify genes that are more specific to normal and cancer states. To highlight these patterns, we calculated the median and standard deviation of the log-transformed distribution of each gene to calculate modified Z-scores. The positive and negative Z-scores may allow us to identify genes that change significantly in expression between cancer and normal states.

### Proteomics analyses

Immunohistochemistry staining for proteins is derived from the Human Protein Atlas (PA) (30). The PA provides immunohistochemistry-based measurement of protein expression for normal and cancer tissue microarrays. The PA’s approach to reporting IHC staining intensity is different for normal tissue and cancer data. For cancer, the PA documents “high”, “medium”, “low”, or “not detected” expression and the number of patients whose tumors fall into those categories for each protein. For normal tissue, the PA documents simply “high”, “medium”, “low”, and “not detected” for each protein in the glandular tissue of the prostate. The normal tissue data does not indicate whether multiple patients were used for testing, and just provides single categorization for protein staining intensity. Essentially, the cancer IHC data is semi-quantitative, the normal IHC data is entirely qualitative, and both are ordinal. Both the cancer and normal tissue specify that expression is assessed in the glandular tissue, improving comparability because prostate cancer cells are glandular (epithelial) in origin.

There are other particularities in the PA data relevant to this work: Staging information for the proteomics data is not provided: It is uncertain whether samples contributing protein stain scores came from a balanced cohort across stages. Also, there is a reliability score describing confidence in IHC staining for normal samples, but not cancer samples. Reliability assessment involves expert review that integrates existing literature, RNA-seq data, and assessment of the IHC staining patterns. Listed in increasing reliability are *Uncertain, Approved, Supported,* and *Enhanced*. For this analysis, we only considered data whose quality was either *Supported* or *Enhanced*.

The differences in data formats made it necessary to transform and normalize data before visualization. For the cancer sample, “High”, “medium”, “low”, and “not detected” were replaced by values of 3, 2, 1, and 0 respectively. The number of patients in each category were then multiplied by these values. For example, if 10 prostate cancer patients had “high” IHC staining intensities, the new calculated values for protein expression would be 30. If five patients had “high” and five had “medium”, the protein expression would be 25 (5 * 3 + 5 * 2). This final calculated cancer protein expression score is therefore a composite of the number of patients in which the protein was detected and the strength of protein staining. Because the cancer samples had different numbers of patients contributing to IHC scores for each protein, we normalize the data by dividing the final IHC score by the number of patients. The normal tissue IHC data was not transformed and remained as an ordinal variable.

### Baseline comparison of Scissor and DEGAS

DEGAS is capable of learning both classification and Cox regression tasks. To provide a baseline for each of these methods, we analyzed a sample of Ductal Carcinoma in-situ (DCIS) from 10x Genomics. DICS is a precursor to invasive breast carcinoma, as in other epithelial-origin cancers, like prostate cancer. The baseline comparator for DEGAS was Scissor (31). Both algorithms were originally designed for scRNA-seq data. Scissor optimizes the linear, logistic, and cox regression models to predict patient outcomes with a Pearson correlation matrix between cell-level and patient-level RNA expression. In this experiment, both algorithms were given the same gene set as input with increasing amounts of sparsity added. The 3 K-means clustering provided by 10x Genomics, aligning largely with tumor, stroma, and immune regions, were used to find the top 300 most differentially expressed genes for each cluster in the ST slide. Next, using the RNA-seq data from the TCGA-BRCA project, we split the data into two groups, patients with relapse/death before 750 days and those with relapse/death after 1500 days (i.e., relapse-free survival). The median follow-up was 761 days. The top 450 most differentially expressed genes were identified for patients who relapsed/survived less than 750 days and for those who relapsed/survived longer than 1500 days. We used the intersection of these two gene sets as the input feature set to DEGAS and Scissor (146 genes). For other experiments, as described earlier, we train DEGAS several times with increasingly large gene sets to provide a regularized output. We do not do that in this experiment because Scissor is not meant to be used in that same way. The relapse free survival (RFS)-derived risk from the TCGA-BRCA data was overlaid onto the cells from DEGAS-BlankCox, DEGAS-ClassCox, and Scissor. For Scissor, the α term was set to 0.0 to force a clinical phenotype correlation output for every spot. Because Cluster 2 from the 10x Visium data almost exactly matches the tumor regions from histology, we measured the alignment of DEGAS and Scissor-derived risk scores with Cluster 2 to assess accuracy. The evaluation metrics were receiver operating characteristic area under the curve (ROC-AUC) and precision-recall area under the curve (PR-AUC). We injected incremental sparsity to simulate drop-out events (0%, 50%, 75%, 90%, 95%) and compared the stability of DEGAS and Scissor performance.

### Integration with single-cell data and C-SIDE

Prostate cancer single-cell data was gathered from Song et al (32), who describe similarities in transcriptional profiles between normal and malignant epithelial cells of the prostate. The single cells were annotated as *Erg+ tumor, Erg– tumor, luminal epithelia, basal epithelia, endothelial cells, fibroblasts, T-cells, B-cells, myeloid cells, plasma cells, mast cells, club cells,* and *smooth muscle*. ERG translocations are a common feature of prostate cancers (33). Using the tool *RCTD-Robust cell type decomposition* (34), these identities were mapped to each spot in the seven spatial transcriptomics samples from Erickson et al. Then, the tool *C-SIDE (cell type-specific inference of differential expression)* from the same research group (35) was used to identify cell-specific changes in gene expression associated with the DEGAS-hazard at each location in the ST arrays. We used *Seurat v4* (36) to interface with these packages.

## RESULTS

### Baseline comparison of Scissor and DEGAS

Using our DEGAS framework, we overlaid risk derived from TCGA-BRCA samples with relapse-free survival (RFS) information (**Fig. 2A**). We identified Cluster 2 as coinciding nearly perfectly with the pathologist-annotated invasive carcinoma regions of the ST slide (**Fig. 2B,C**). We found that both methods performed remarkably well on ST data considering neither are designed for use with ST data (**Fig. 2D-L**, **Table 1**). The DEGAS-BlankCox, which does not use ST cluster labels during training, assigned higher risk to the invasive carcinoma regions, and this correlation between risk and invasive carcinoma regions decreased as more sparsity was introduced (**Fig. 2D-F, Table 1**). Likewise, the DEGAS-ClassCox model, which does use clustering labels, performed well across all levels of sparsity though with some decreased performance at the higher levels of sparsity (**Fig. 2G-I**, **Table 1**). Scissor also performed well, especially when no sparsity was added but had marked decreases in performance with increasing sparsity (**Fig. 2J-L**, **Table 1**). Given ST data is already very sparse with many false negative gene detections, it is not surprising that adding more sparsity will drastically reduce performance for any method. It should be noted that Scissor is designed to select a subset of the most extreme high-risk cells/spots based on the distribution of correlation score outputs. It is not intended to give an overall score to every cell/spot. In this context Scissor is not addressing the exact same problem as DEGAS. But, overall, both methods are viable for use with ST data. DEGAS without training on ST cluster labels (DEGAS-BlankCox) performed well and adding cluster labels during training (DEGAS-ClassCox) made the model more resilient to drop-out events (**Fig. 2G-I**, **Table 1**).

**Figure 2:**
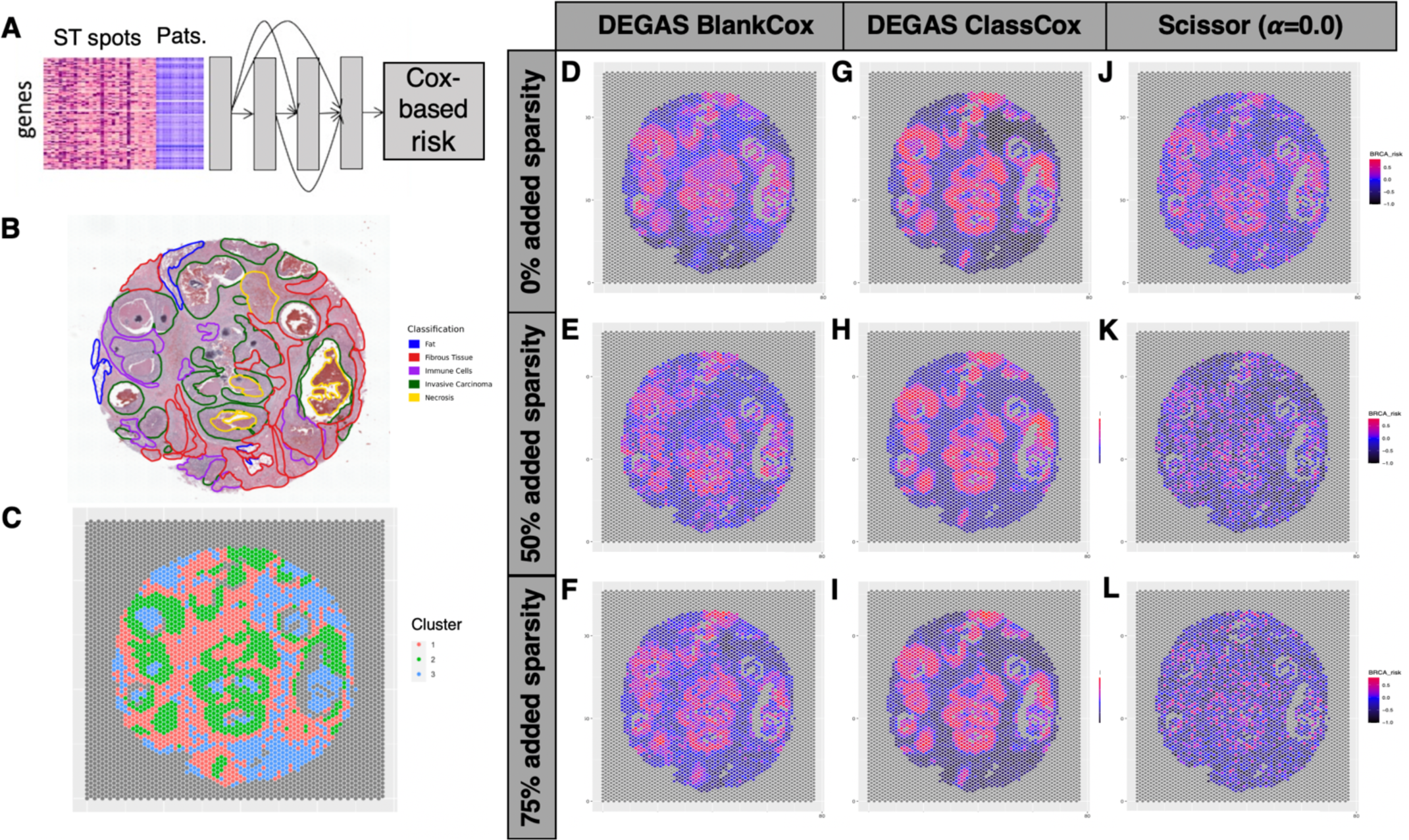
Baseline comparison of DEGAS and Scissor in predicting Cox hazards on data of increasing sparsity. Baseline comparisons for DEGAS models and Scissor. **A.** Overview of DEGAS prediction. **B.** Pathologist labels for ST slide. **C.** Cluster labels from 10XBRCA1 ST slide. **D-F.** DEGAS-BlankCox output for different levels of sparsity 0% **(D)**, 50% **(E)**, and 75% **(F)**. **G-I.** DEGAS-ClassCox output for different levels of sparsity 0% **(G)**, 50% **(H)**, and 75% **(I)**. **J-L.** Scissor output for different levels of sparsity 0% **(J)**, 50% **(K)**, and 75% **(L)**.

**Table 1.** Performance metrics, ROC-AUC and PR-AUC, for DEGAS-BlankCox, DEGAS-ClassCox, and Scissor (α=0.0) trained on 10XBRCA1 and TCGA-BRCA datasets.

|  | DEGAS-BlankCox |  | DEGAS-ClassCox |  | Scissor |  |
| --- | --- | --- | --- | --- | --- | --- |
| Sparsity | ROC-AUC | PR-AUC | ROC-AUC | PR-AUC | ROC-AUC | PR-AUC |
| 0% | 0.9143 | 0.7462 | 0.9906 | 0.9785 | 0.8897 | 0.7395 |
| 50% | 0.7967 | 0.5559 | 0.9843 | 0.9542 | 0.7566 | 0.5622 |
| 75% | 0.9091 | 0.7992 | 0.9792 | 0.9467 | 0.6648 | 0.4599 |
| 90% | 0.7764 | 0.6267 | 0.9593 | 0.9060 |  |  |
| 95% | 0.6459 | 0.4358 | 0.9284 | 0.8562 |  |  |

### DEGAS risk impressions in normal prostate, Stage II, Stage III, and Stage IV adenocarcinomas

DEGAS highlights areas of glandular tissue in the normal prostate sample as being associated with poor progression-free survival (i.e., *“High risk”*, Fig. 3A). The sample was obtained from the public 10x repository, with no metadata as to the demographics of medical history of the patient. Pathologists review of the H&E stained whole-slide image confirmed that the heat map was highlighting glandular tissue, and that the highest-hazard region **(Fig. 3A black arrow)** is a gland with no morphology particularly suspicious for malignancy, nor are the other, low-risk glands in this specimen **(Supp. Fig. 3A)**. Specifically, there was no evidence of neoplasia, noting only that there may be some urothelial metaplasia, but this is a benign process where urothelial cells can intermix with normal prostatic glands and ducts (37). It is not cancer and is not atypical. Histologic correlates of the Stage II prostate adenocarcinoma high risk regions were difficult to determine because there was no diagnostic H&E-stained image associated with the sample (**Fig. 3B**). In the Stage III prostate adenocarcinoma (**Fig. 3C**), there is a high-risk region (red, left), which is the bulk cancer tissue. On the right is the low-risk peritumoral stroma and normal prostatic glands (blue), within which a solitary nerve is highlighted as high-risk (black arrow). In the Stage IV sample (**Fig 4D**), the high-risk regions coincide with slightly higher Gleason score morphologies, and the low-risk regions had more normal glandular tissues. Highest risk region shows back-to-back glands a high concentration of lymphocytes (**Supp. Fig. 3B**). Histological labels for the normal tissue are provided in **Fig. 4E**. Intuitively, DEGAS risk impression increases with stage (**Fig. 4F**). Based on risk output by DEGAS, we perform differential expression on the high-risk regions of each sample (**Fig. 4G**). Differential expression analysis in the normal sample would have been biased if we used solely the risk labels to define high and low risk: stromal regions are generally low-risk and we do not want to compare stroma against glands. Therefore, we used histological labels to compare only high and low-risk glands and removed stroma from the normal tissue analysis. UMAP and t-SNE of the top 3000 most variable genes among all four samples shows high-dimensional colocalization of ST spots: The Stage IV sample has a distinct cluster, and the highest risk regions of normal tissue localize to this cluster in the UMAP (**Supp. Fig. 4**). DEGAS aligns the distribution of genes between patients and ST samples in latent space using the maximum mean discrepancy loss (MMD). Colocalization of ST spots in UMAP and t-SNE derived embeddings provides some intuition to the predicted risk scores for a few of the most high-risk ST spots, but do not fully explain the continuous output from DEGAS for all spots that clearly demarcate histologic regions.

**Figure 3:**
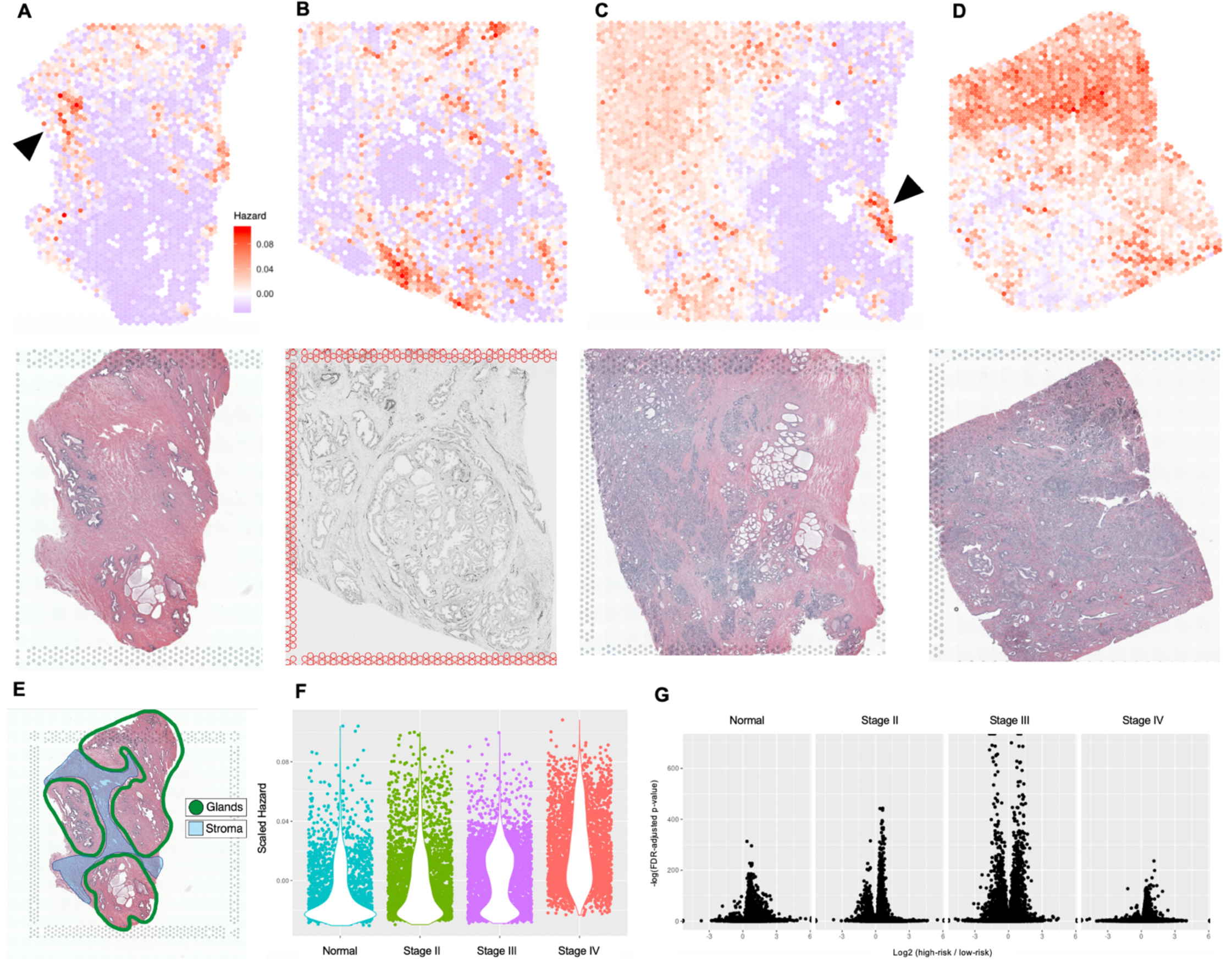
DEGAS risk impressions in four spatial transcriptomic samples of prostate tissue. Hazard predictions for four prostate ST samples. High (red) and low (blue) risk scores from the DEGAS model are visualized. All samples share the same hazard/color scale. **A.** Histologically normal prostate tissue, with black arrow indicated a particularly high-risk region, with corresponding H&E-stained section below. **B.** Stage II prostate adenocarcinoma with tissue section below. This specimen was stained with immune-fluorescence and thus not H&E-stained. **C.** Stage III prostate adenocarcinoma with right-adjacent normal tissue (blue), and an arrow indicating a particularly high-risk region. In the corresponding H&E, below, examination by a pathologist revealed that the region is a nerve. **D.** Stage IV prostate acinar cell carcinoma (i.e., adenocarcinoma) and corresponding H&E section below. **E.** Histological labels from normal prostate tissue shown in A. **F.** Scatter plot overlaid with violin plot of the hazards for all ST spots in Panels A-D, showing trend of increasing hazard with increasing stage. **G.** Differential gene expression plot using Wilcoxon test FDR-adjusted p-values and the log 2 mean fold change the samples in panels A-D.

**Figure 4:**
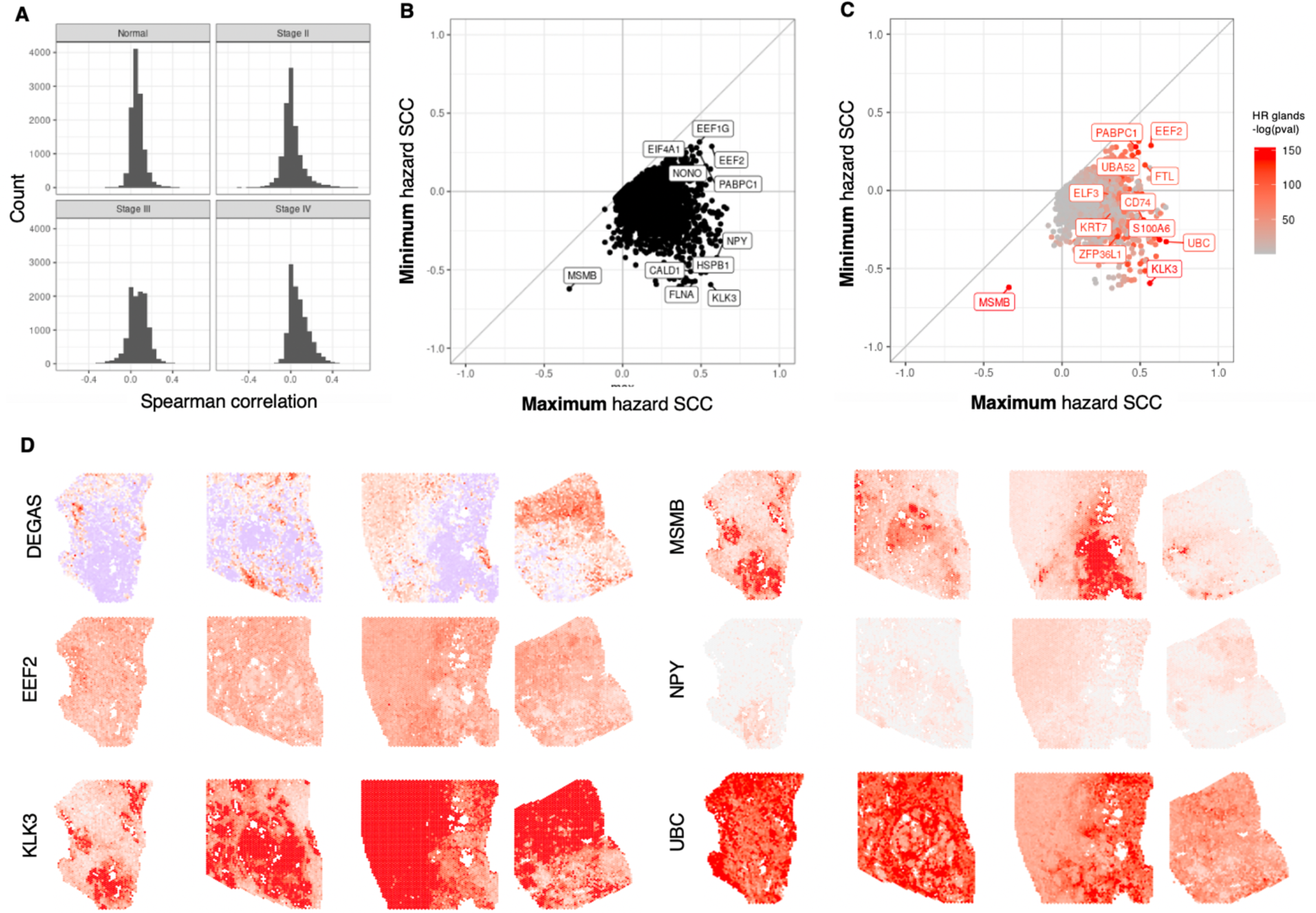
Coefficients of Spearman correlation between genes and DEGAS hazard output across all ST spots on all prostate cancer samples. **A.** Histogram of correlation coefficients between genes and the DEGAS hazard output for each spot. The distributions are different. For the normal sample, the Spearman correlation was not restricted to only glandular tissues (thus, the title refers to “all ST spots”). **B.** Concordance between Spearman correlation coefficients (SCC) across the different prostate samples is demonstrated by plotting the range of coefficients across the four samples (maximum and minimum). MSMB is the only gene for which the spearman correlation is consistently negative across all slides. Rack1, EEF2, and others are consistently associated with a high hazard. Some genes with positive and negative coefficients are KLK3, NPY, HSPB1, and CALD1. **C.** Panel B overlaid with the FDR-adjusted –log(p-value) for each gene’s enrichment in the high-risk glandular tissues. Labeled genes have –log(p-value) > 120. **D.** Expression of genes from panels B & C visualized on four prostate ST samples, in order from left to right: Normal, Stage II, Stage III, and Stage IV. Degas hazard provided top left for reference, where blue indicates hazard impression below the median value, and red is above the median. KLK3 is expressed in both normal glands and cancer tissues, but not in the highest risk glandular region. MSMB is expressed in normal tissues, but not in cancer, and not in the highest risk normal glandular region. NPY is expressed in glandular tissue, as well as some of the cancer tissues, but not in the highest risk normal glandular region. UBC is somewhat opposite to the NPY pattern, highly expressed in multiple normal regions, including the high-risk glandular region, diffusely in the Stage II cancer, mostly in the normal regions of the Stage III cancer, and diffusely in the Stage IV cancer.

### Differential expression in high-risk normal glandular region

We next investigated functional enrichment of differentially expressed genes among the high and low risk glands in the normal sample. Enrichment for *Disease* ontologies include *Lupus Erythematosus, Systemic* (P = 2.86×10^-9^), *Triple Negative Breast Neoplasms* (P = 6.53×10^-9^), *Anophthalmia and Pulmonary Hypoplasia* (P = 8.68×10^-9^) and *Neoplasm invasiveness* (P = 9.91×10^-9^). Top terms in the *GO: Biological Process* category are *Cell-cell adhesion* (P = 2.79×10^-7^), *Epithelium development* (P = 7.29×10^-7^), *Antigen processing and presentation of exogenous antigen via MHC Class II* (P = 8.81×10^-6^), and *Leukocyte cell-cell adhesion* (P = 1.02×10^-5^). Enrichments in the *Cellular Component* ontology term include *Lysosome* (P = 2.07×10^-12^), *Focal adhesion* (P = 2.37×10^-7^), *Secretory granule* (P = 5.3×10^-7^), and *MHC Class II protein complex* (P = 7.98×10^-7^). *Molecular Function* ontologies include *MHC Class II binding* (P = 7.11×10^-6^) and *cell adhesion molecule binding* (P = 1.74×10^-6^). *Interactions* are enriched for *AGR2* (P = 8.84×10^-14^), *FN1*, *BTF3*, *CYLD*, *ITGA4* (all P = 1.83×10^-12^), *TNF* (P = 5.67×10^-12^), and *FOLR1* (P = 3.20×10^-8^).

Four enrichment categories overlapped between the high-risk regions of the glandular normal tissues, Stage II, Stage III, and Stage IV samples: *High-grade prostatic intraepithelial neoplasia*, *Hormone refractory prostate cancer, Prostatic neoplasms,* and *Renal Cell Carcinoma* (**Table 2, Supp. Fig. 4**). The genes contributing to the PIN module are not the same for each sample. The lack of KLK3 enrichment in the high-risk normal sample is conspicuous, given that KLK3 (PSA) is the main screening marker for prostate cancer available to clinicians. The expression of the genes for the normal samples are visualized (**Supp. Fig. 6**): Notably, β-Catenin 1 (CTTNB) has lower expression in the low-risk glandular region and higher expression in both the stroma and highest-risk glands.

**Table 2:** Overlap in functional enrichment among High-risk Normal glandular prostatic tissue, and Stage II, III, IV samples

| Category | Enrichment term | Genes from high-risk normal glands contributing to enrichment | FDR BH for high-risk normal tissue | FDR BH for high-risk Stage II tissue | FDR BH for high-risk Stage III tissue | FDR BH for high-risk Stage IV tissue |
| --- | --- | --- | --- | --- | --- | --- |
| Disease | High-Grade Prostatic Intraepithelial Neoplasia | ACTB, GSTP1, IGFBP3, SQSTM1, CTNNB1 | 3.78E-03 | 2.23E-03 | 4.32E-03 | 9.49E-03 |
|  | Hormone refractory prostate cancer | PSAP, HMGB1, PABPC1, DDX5, GSTP1, IGFBP3, CLU, AHNAK, SLPI, CTNNB1, MMP7, NONO, HSPB1 | 9.46E-04 | 1.65E-04 | 1.74E-03 | 3.05E-04 |
|  | Prostatic Neoplasms | LITAF, B2M, RUNX1, GSTP1, IGFBP3, ZFP36L2, CLU, EHF, CTNNB1, CTSB, CTSD, CNN3 | 1.40E-03 | 3.47E-05 | 5.61E-06 | 3.05E-04 |
|  | Renal Cell Carcinoma | ACTB, GSTP1, IGFBP3, SQSTM1, CTNNB1 | 2.84E-07 | 2.16E-11 | 8.37E-03 | 4.91E-05 |

### Correlation of DEGAS hazards and gene expression

The prior experiments demonstrated that high-risk tissues are not enriched for the same genes and ontologies. To identify genes that were consistently correlated with DEGAS risk scores, we calculated the Spearman Correlation Coefficients (SCC) between DEGAS Hazards and each gene’s expression within each of the four samples. By calculating the SCC for each sample independently, we described the range of correlations across all four samples. Plotting the minimum and maximum SCC’s allowed for quick identification of consistently correlated genes. The distribution of correlation coefficients is different for each sample **(Fig. 4A)**. MSMB is the only gene strongly and consistently negatively correlated with high hazard regions **(Fig. 4B)**. This gene is lowly expressed in the highest-risk region of normal glandular tissue. Several genes like *EEF2*, *PABPC1*, *FTL*, and *UBA52* are consistently positively correlated with high hazard regions. Many genes have inconsistent correlations, like *KLK3* (PSA), *S100A6*, *UBC*, and *ZFP36L1* which are negatively and positively correlated in different samples **(Fig. 4C)**.

Visualizing the gene expression of these archetypes reveals distinct spatial patterns **(Fig 4D)**: *EEF2* is correlated with hazard, but with low specificity to glandular or stromal tissues. *KLK3* is strongly expressed by both cancer and glandular tissue but is down-regulated in the highest-risk glandular tissue. *MSMB* is also downregulated in the highest-risk glands but, unlike *KLK3*, its expression is restricted to non-cancerous epithelial cells. *NPY* is expressed in the low-risk, but not high-risk normal glandular tissue. *NPY* appears in bulk cancer region of the Stage III and Stage IV cancers. *UBC* is expressed in multiple regions of the normal tissue, especially the high-risk region, diffusely in the Stage II sample, opposite the risk scores in the Stage III sample, and diffusely in the Stage IV sample. DEGAS hazard is tightly correlated with total RNA expression (SCC 0.546, **Supp. Fig. 7**).

### Integration of proteomics and RNA-seq

To understand how genes differentially expressed in the ST data were represented in large cohorts, we integrated RNA-seq from patients with cancer (TCGA) and normal tissue form healthy donors (GTEX). First, we noticed that overall, the median gene expression well-correlated between cancer and normal tissues for most genes, even for cancer-hallmark genes like *TP53*, *PTEN*, and *APC* **(Fig. 5A)**. Similarly, in the Human Protein Atlas IHC data, there was a general trend where proteins that were more robustly expressed in normal tissue were also increasingly found in cancer tissues (**Fig. 5B**). Integrating RNA-seq and proteomics in the showed the same trend, where genes with increasing RNA expression tended to be more highly expressed at the protein level in both cancer and normal tissues (**Fig. 5C**).

**Figure 5:**
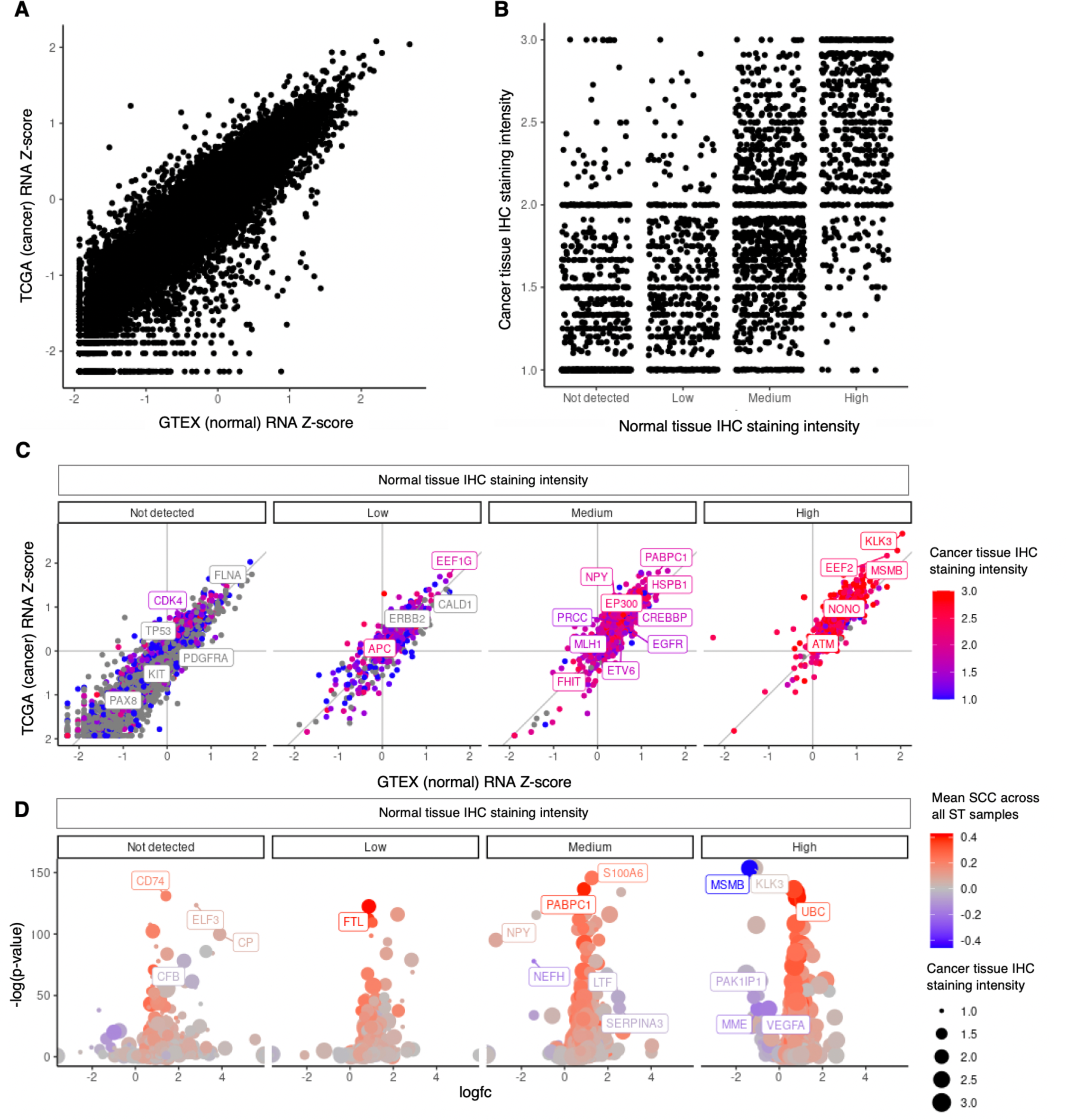
RNA expression z-scores and protein abundance across tumor and normal tissues. Bulk sequencing do not demonstrate nuance of DEGAS-identified ST data, though DEGAS was trained using this lower resolution RNA-seq. **A.** Modified z-scores of RNA expression show that tumor (TCGA) and normal (GTEX) are highly correlated with one another. **B.** There is also a general correlation of protein expression between cancer and normal tissue, where higher RNA z-scores in both cancer and normal are associated with higher protein expression in both higher and normal. **C.** As RNA expression increases, protein expression in both cancer and normal tissues increases. *KLK3* is highly expressed in both cancer and normal tissues in this low-resolution scale. **D.** Volcano plot comparing RNA expression in the high-risk regions among the glandular tissue in the normal ST sample. Color corresponds to the Spearman Correlation Coefficient averaged across all four samples. Size of points reflects the calculated score for IHC staining intensity for proteins in cancer.

Overlaying the proteomics on the differential expression of normal prostate high and low risk regions revealed several patterns (**Fig. 5D, Supp. Fig. 8**): Proteins like *CD74*, *CP*, *CFB* and *FTL* are not strongly expressed in normal tissue, more highly expressed in cancer, and the RNA are highly upregulated in the high-risk normal glands. *FTL* is also positively correlated with DEGAS risk across all four samples (SCC in normal sample = 0.470, Stage II = 0.531, Stage III = 0.162, Stage IV = 0.468). Both *NEFH* and S100A6 are moderately expressed in normal tissue, but less so in cancer tissue. *NEFH* is downregulated in the high-risk normal region and *S100A6* is upregulated. Both have moderate correlations with DEGAS hazards. *NEFH* is an intermediate filament that helps maintain the structure of neuron axons and soma (38). *S100A6* is a cell-cycle protein whose up-regulation is associated with keratinocyte de-differentiation and proliferation (39). As stated previously, prostate glands are epithelial tissue. Again, the Protein Atlas data demonstrates that *KLK3* and *MSMB* can be expressed in both normal and cancer tissues. DEGAS has highlighted a nuance, beyond the bulk data, where the highest-risk glands, which shows signatures of carcinogenesis, do not express these proteins.

#### Identification of high-risk benign morphology glands in validation dataset

To identify consistent and reproducible genetic findings, we analyzed an additional prostate cancer spatial transcriptomics dataset. This particular data has recently been used to demonstrate that copy number variations (CNV’s) occur in both benign and malignant tissues (15). The authors provide compelling evidence for epigenetic changes in early carcinogenesis, and shared lineage among clones in both benign and cancerous tissue, aligning with the multifocal origins of prostate cancer in 60-90% of cases (9). One morphologically benign sub-clone in sample H2_1, Clone C, had a gain of MYC activity, loss of *MSMB*, and downregulation of *KLK2*, *KLK3*, *FKBP5*, *NKX3-1*. The latter are androgen-associated genes (40) and overlap with our prior findings. Per the original publication, this clone C also demonstrated upregulated in transcriptional signatures consistent with phenotypic versatility.

Applying DEGAS to this important dataset showed a correlation between the Hazard predictions and CNV in the morphologically benign glandular tissues (SCC = 0.652), and again highlighted regions of normal glandular tissue that were high risk **(Fig. 6AB)**. As in prior experiments, there is an intuition with DEGAS output: hazard tends to increase with increasing tumor grade **(Fig. 6C)**, histological labels of *Prostatic intraepithelial neoplasia* and *Transition state* (denoting tumor/normal cell mixtures at interfaces) received higher risk scores, but the normal prostatic epithelium had a large variation in DEGAS hazards. Closer inspection of the normal sample scores showed a multi-modal distribution **(Fig. 6D)**. Plotting these four modes revealed distinct patterns in the tissue, with *Benign gland* (BG) *risk rank 1* (lowest risk) rarely overlapping with *BG risk rank 4* (highest risk). BG Risk ranks 3 and 4 have intermediate risk and show more colocalization.

**Fig. 6:**
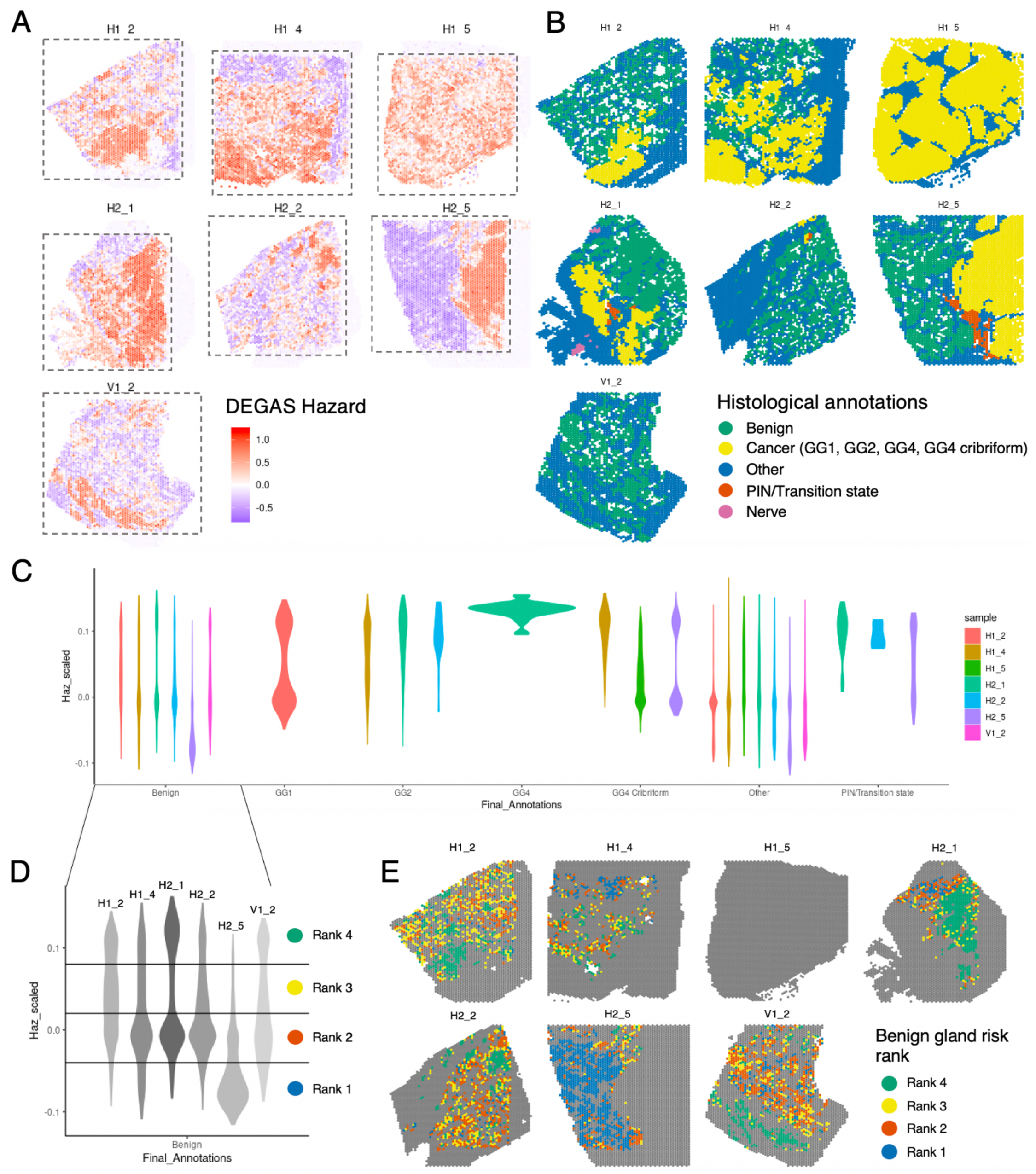
DEGAS highlights high risk morphologically benign glands with discrete spatial localization in external dataset. **A.** DEGAS hazard impressions for seven Visium ST samples from the prostate of a single patient. **B.** Across these seven samples, there is a variety of both cancerous and benign histology. **C.** Violin plots of DEGAS hazards shows a trend between increasing tumor grade and hazard predictions across GG1 through GG4 regions. **D.** The morphologically benign tissues have multi-modal DEGAS Hazard distributions and can be divided into four ranks of increasing hazard. Visualizing these four ranks reveals discrete spatial distributions. Spatial coordinates are not used in DEGAS training and prediction.

Differential gene expression of these risk groups shows **(Fig 7)** that several genes transition from being highly to lowly expressed (and *vice versa*) across the BG risk groups. There number of ST spots in each rank is as follows: Rank 1, 1655 spots; Rank 2, 1039 spots; Rank 3, 1402 spots; Rank 4, 882 spots. BG rank 1 and 4 have the most differential expression among groups. *MSMB* and *NEFH* are highly expressed in BG rank 1 (low-risk) and transition to low expression in BG rank 4 (highest-risk). *RPL11* and *OR51E2* are lowly expressed in low-risk tissue, but transition to high expression in the high-risk tissue (Fig 7AB). This data can be visualized as a pseudo-trajectory along DEGAS-defined risk groups (Panel C).

**Fig. 7:**
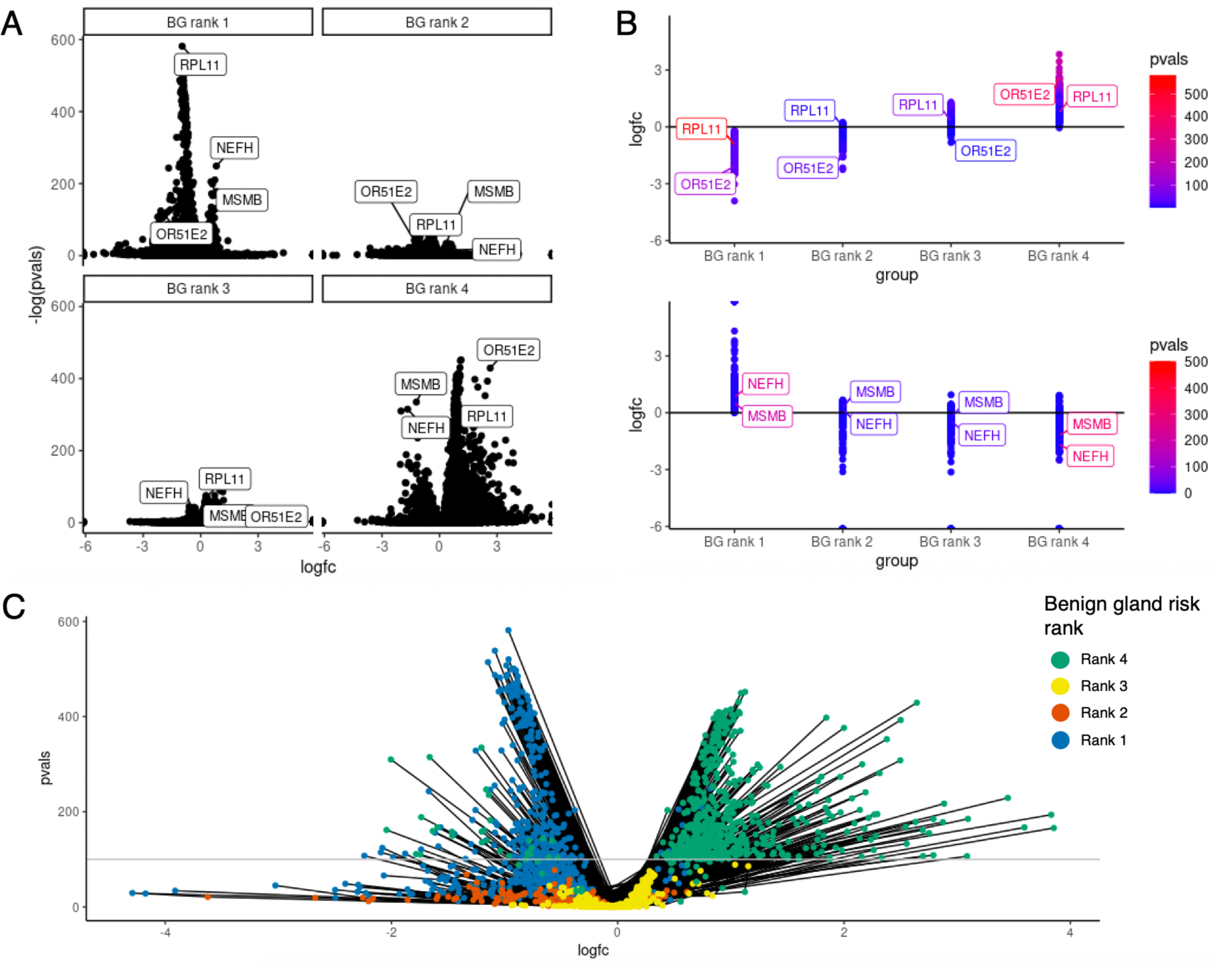
Differential expression of morphologically benign gland risk ranks highlights gene trajectory in transition from low risk to high-risk cell states. **A.** Wilcoxon differential expression of Benign Gland (BG) risk ranks, where Rank 1 is the lowest risk, and Rank 4 is the highest risk. This comparison is performed by comparing each rank to the pooled expression of all other ranks, with FDR-adjusted p-values calculated on the concatenated four comparisons. BG Rank 1 and Rank 4 have the most polarized expression. Certain genes, like *MSMB* and *NEFH* are highly expressed in lower-risk Rank 1 and are suppressed in the higher-risk Rank 4. *OR51E2* has the opposite pattern, with lower expression in normal tissue and higher expression in high-risk tissue **B.** Plotting the log fold-change of genes, and using the risk rank as a pseudo-time, shows that some genes, like *MSMB* and *NEFH* have a consistent progression from low to high expression. **C.** Combining the genes with -log(p- values)>100 in any of the four risk ranks into a single volcano plot, shows that many of these highly significant genes transition from low to high expression from low to high-risk morphologically benign glands. Lines track the progression of a single gene across each rank, demonstrating that Ranks 2 & 3 have fewer distinct genes than Ranks 1 & 4.

ST spots harbor multiple cells within their diameter. We were therefore interested in deconvoluting/decomposing cell identities within each ST spot and understanding how cell-specific gene expression changes with DEGAS risk across these tissues. The tool RCTD was used to transfer annotated cells from a recently-published single-cell analysis of prostate cancer onto the spatial transcriptomics **(Fig. 8)**. RCTD provides cell-type predictions as either doublets or singlets, where doublets have two distinct cell types predicted at each location (e.g., Club cell and fibroblast) and singlets are where only a single cell type was attributed to the location. Bulk regions of tumor (e.g., sample H1_5) contained large populations of *ERG* + tumor cells **(Fig. 8A)**. *ERG* – tumor cells were attributed to many of the histologically normal glands **(Fig. 8B)**. Luminal and basal epithelial cells were also attributed to the high-risk normal gland regions identified by DEGAS **(Fig. 8C)**. Plasma and myeloid cells were widely prevalent in the two samples, and there is a small group of B-cells in the bottom- right of sample V1_2 **(Fig. 8D)**. This nidus corresponds to what appears to be a tertiary lymphoid structure **(Supp. Fig. 9)**, aligning with increased *CD19*, *CD27*, and *CXCL13* expression (41).

**Fig. 8:**
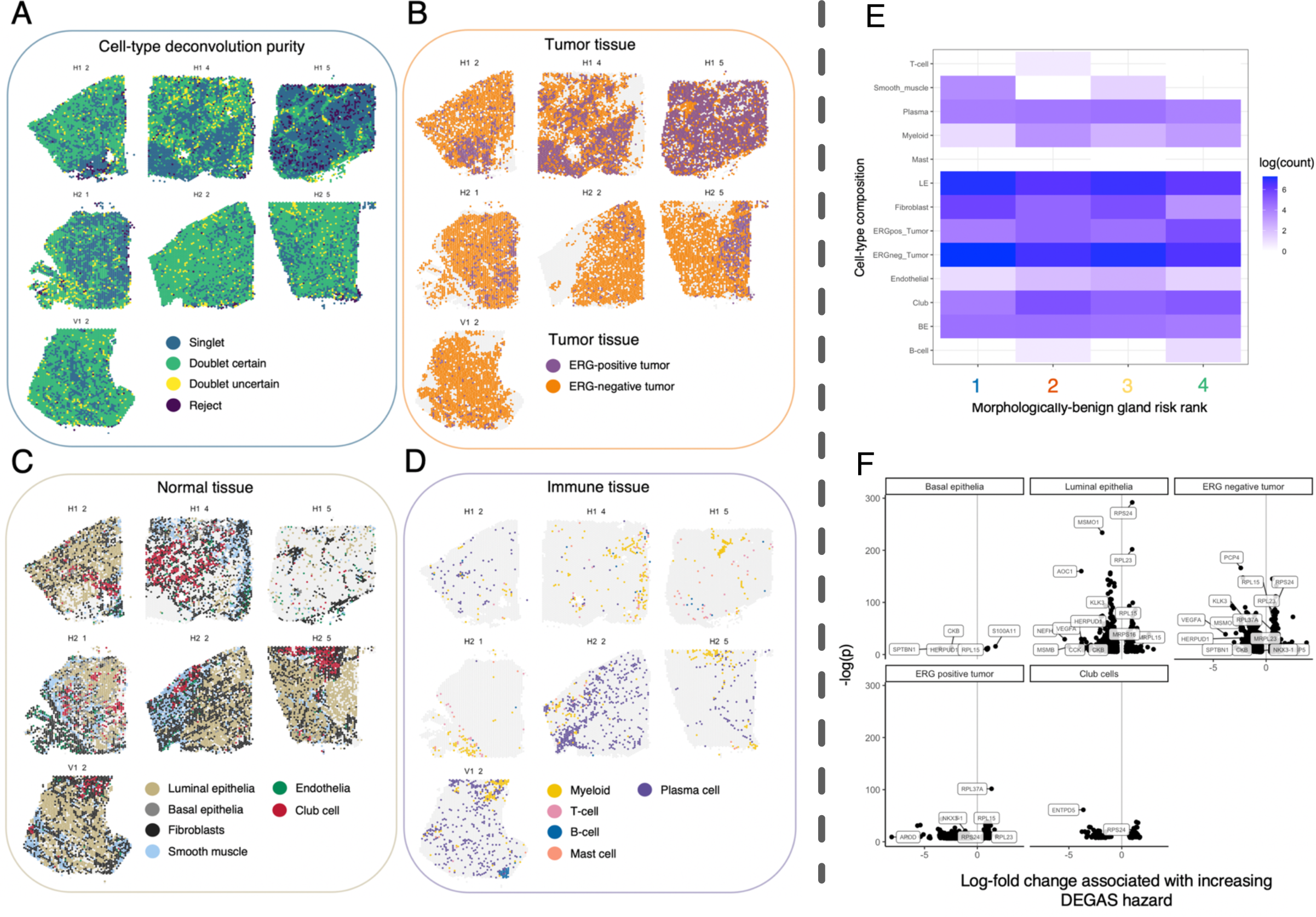
ST-spot deconvolution with annotated single-cell dataset and cell-specific differential expression associated with DEGAS risk. **A.** Cell deconvolution/decomposition at each ST spot by the RCTD attributes single cells to each ST spot. If there are multiple predominating cell types at a location, RCTD provides “doublets”. Certain locations cannot be confidently decomposed and are called “reject”. H1_5 is the majority cancer sample and is attributed mostly a single cell type. Other samples show greater heterogeneity in the number of cell-types ascribed to each location. **B.** Both ERG-positive and negative tumor cells are mapped both tumor and non-tumor histological regions. **C.** Normal tissue contains predominantly luminal epithelia, and tumors also have luminal cells. **D.** Immune cells, mostly myeloid and plasma cells, are scattered throughout each sample. There is a group of Bcells in the lower right corner of the V1_2 sample. **E.** Cell-type composition of the four morphologically-benign gland risk ranks is largely consistent. **F.** Cell-type specific differential expression associated with DEGAS.

Cell-type composition does not vary widely among the BG risk ranks **(Fig. 8E)**, implying a cellular state change rather than a composition change. The BG regions contain mostly luminal epithelial and *ERG* – cancer cells. The largest differential expression comes from luminal epithelial cells and *ERG* – tumor regions **(Fig. 8F)**. Cell-type specific differential expression associated with DEGAS risk scores show that the *MSMB* and *NEFH* downregulation is specific to high-risk luminal epithelial cells. *MSMO1* (*SC4MOL*) is negatively associated with DEGAS risk in luminal epithelial cells. Basal epithelia can be identified by expression of *S100A11* (42), and C- SIDE analysis revealed that this gene was associated with higher DEGAS risk scores. The basal cells also show downregulation of *CKB*. Many genes positively associated with risk in the *ERG* –/+ tumor tissues are ribosome- associated proteins (e.g., *RPL37A*, *RPL23*, *RPS24*).

## DISCUSSION

### Baseline comparison

Both DEGAS and Scissor perform well with spatial transcriptomics. At very high sparsity levels, DEGAS outperforms Scissor, with ROC-AUC 0.901/0.9792 vs. 0.6648 at 75% sparsity respectively. It should be noted Scissor was not designed to output a prediction for every single point, just those at the extreme ends of correlation with clinical outcomes. The baseline comparison showed how DEGAS trained with cluster labels for the ST data provides an output smoothing that is more resistant to dropout events, though the model without class labels (*BlankCox*) also had stable performance across all levels of sparsity. Scissor consistently highlighted high risk cells which coincided with the tumor regions, which was expected and consistent with DEGAS. Both Scissor and DEGAS performed very well, considering neither technique was designed specifically for ST.

### Risk impressions, morphological and functional enrichment interpretation

In the normal tissue, there were benign morphology glands whose transcriptomic signatures were associated with cancer and carcinogenesis, including *Triple Negative Breast Neoplasms* (P = 6.53×10^-9^) and *Neoplasm invasiveness* (P = 9.914×10^-9^). Enrichment for *Cell-cell adhesion* (P = 2.79×10^-7^), *Epithelium development* (7.29×10^-7^), *Antigen processing and presentation of exogenous antigen via MHC Class II* (P = 8.81×10^-6^), and the *Lysosome* (P = 2.07×10^-12^) are also seen (**Fig. 3AE, Supp. Fig. 1**). Being upregulated processes, this may represent tissue that is being engaged by the patrolling immune system via HLA pathways. Interestingly, the enrichment refers to MHC Class II. Traditionally, MHC Class I are viewed as the mediators of endogenous protein antigen presentation; however, inflammatory signals can spur non-professional antigen presenting cells to express MHC Class II and present endogenous antigens via lysosomal pathways (43).

The high-risk normal glands are also enriched for interactions with *AGR2* (P = 8.84×10^-14^), *FN1*, *BTF3*, *CYLD*, *ITGA4* (all P = 1.83×10^-12^), *TNF* (P = 5.67×10^-12^), and *FOLR1* (P = 3.20×10^-8^). Folate is involved in DNA synthesis, and therefore, it’s logical that *FOLR1* is linked to cancer progression across several cancers, *Wnt/ERK* signaling, and commonly overexpressed epithelial cancers (44). This has made it a promising resource for targeted drug-delivery (45), and slight upregulation of *FOLR1* in high-risk benign tissues was seen in Erickson et al. dataset (P = 1.2×10^-4^, log2FC = 1.31). *FN1* is a glycoprotein that connects the cell to supporting stroma and involved in cell migration. *FN1* is known to be involved in the epithelial-to-mesenchyme transition (EMT). Normal tissue is thought to express this protein at lower levels than cancer cells; the hypothesis being that cancer cells increase expression to favor cell-stroma interactions, rather than the cell-cell interactions which normally inhibit continued cell growth. *FN1* was not significantly upregulated in the BG risk group 4.

Histologic correlates in the Stage II sample could not be determined, because the specimen was not H&E-stained, so comments on morphology herein do not pertain to the Stage II sample (**Fig. 3B)**. The Stage III sample showed the bulk tumor as being higher risk than the stroma, which makes intuitive sense. Stromal tissues were most low risk, except for a solitary nerve, which was highlighted as being high-risk (**Fig. 3C**). The Stage IV cancer highest risk regions were glands with higher Gleason scores, with dense glands, lymphocytic infiltration, and some free cancer cells that do not obey the original glandular architecture (**Fig. 3D, Supp. Fig. 3**). This is intuitive, as higher Gleason grades are associated with a worse prognosis (46).

The presence of this high-risk nerve in the Stage III sample and *NEFH* expression in normal tissue (a neuron cytoskeletal protein) align with existing literature. Parasympathetic and Sympathetic autonomic nerve fibers have been shown contribute to the development and progression of prostate cancer in mouse models: Specifically, Type 1 muscarinic receptors (parasympathetic signaling) are associated with an increased tumor invasion and metastasis, and Type 2 & 3 β-adrenergic receptors in the microenvironment are associated with the initiation and progression of carcinogenesis (47). Cancer spread along nerves (termed *perineural invasion*) is an independent histopathological poor prognostic factor in prostate cancer (48). The parasympathetic nerves are closely associated with the glandular epithelium of the prostate (49). This has made autonomic drugs, specifically beta-blockers an area of interest in the reduction of prostate cancer incidence and recurrence, and the subject of three clinical trials (50), studying propranolol (NCT01857817, NCT03152786) and carvedilol (NCT02944201). Overlap in functional enrichment of high-risk tissues from all four of these tissues include *High-grade prostatic intraepithelial neoplasia*, *Hormone refractory prostate cancer, Prostatic neoplasms,* and *Renal Cell Carcinoma*. Signatures of intraepithelial neoplasia and invasive cancer are surprising to find in a histologically normal prostate. The genes in the high-risk normal tissue contributing to the PIN enrichment are *ACTB*, *GSTP1*, *IGFBP3*, *SQSTM1*, and *CTNNB1*. *CTNNB1* (β-catenin) is involved in the epithelial-to-mesenchyme transition through interactions with cell polarity gene E-cadherin and matrix metalloproteases. This is further explored in the discussion of MSMB.

### Correlation of DEGAS risk scores with gene expression

Correlating the RNA expression and DEGAS risk scores highlighted *KLK3* and *MSMB* as unique proteins (**Fig. 4BC**). *KLK3* is used clinically as a blood test to screen for prostate cancer, track the progression and responses to treatment, and monitor for recurrence. Both *MSMB* and *KLK3* are proteins secreted by normal prostate glands. DEGAS allowed us to detect a nuance: *KLK3* RNA is expressed highly in low-risk normal glands and cancer tissue but is lost in the highest-risk normal gland region. Similarly, *MSMB* RNA expression is lost in the highest-risk regions, but unlike *KLK3*, it is consistently correlated with lower hazards across the normal, Stage II, III, and IV samples. *MSMB* is therefore mostly expressed in low-risk glands and expressed at a lower level in cancer tissues generally. In some of the highest-risk Stage IV regions, it is not expressed. Since *MSMB* secretion is a normal function of differentiated prostate glands, loss of this ability may be a sign of de-differentiation, and loss of *MSMB* may be a specific marker for premalignant lesions (51), prostate cancer generally (52), and is especially reduced in higher grade tumors (53). We noted that the high-risk glands were highly enriched for *S100A6*. Keratinocytes overexpressing this gene in 2D and 3D cell cultures have shown proliferate more quickly, have a greater affinity for fibronectin, and lose protein markers of differentiated keratinocytes, which are another type of epithelial cell (39). Fibronectin was previously discussed for its role in the EMT.

Immunohistochemical quantification of MSMB in 3268 microarrays of prostatectomy specimen has revealed that patients with higher *MSMB* expression have a decreased prostate cancer recurrence after surgical removal of the prostate (54). Single-nucleotide polymorphisms strongly associated with prostate cancer susceptibility lie upstream of *MSMB*: The rs10993994 variant has a strong association with prostate cancer, and in a cell-line assay showed preferential binding to the *CREB* transcription factor which led to lower expression of *MSMB* (55). *CREB* itself is widely involved in cancer and carcinogenesis, being activated by the several phosphorylation cascades, including the *Ras/Raf/MAPK/ERK* pathway (which are downstream of *KRAS*), and has diverse roles in survival, proliferation, and differentiation described across several cancers (56). In pancreatic adenocarcinoma specifically, *KRAS* is a centrally important driver and *CREB* is hypothesized to play a role in the autocrine via *TGF-β* signaling (57). *CREB* is moderately enriched in the high-risk normal glandular tissue analyzed herein (P = 2.19×10^-5^, log2FC = 1.05) as is *TGF-β* (P = 1.05×10^-7^, log2FC = 1.07). *TGF-β* itself is plays a central role in the EMT program used in both normal settings, as in wound healing and fibrosis, as well as in cancer (58).

*MSMB* has been studied with in the prostate using fluorescence in-situ hybridization data (51), showing that: RNA expression be lost in low and high-grade intraepithelial neoplasia, expression is higher in more-differentiated carcinomas, and though prostate acid phosphatase (*PAP*) and *PSA* are expressed by 6-7 weeks fetal prostate tissue, *MSMB* is not yet expressed. Fetal tissue is interesting because the prostate is not fully differentiated until puberty, and the authors note that the pubertal prostate has a variable *MSMB* staining that decreases from the periphery to the central zones. Interestingly, the authors note that *MSMB* staining is stronger in the periphery of the prostate in adults as well, which is where most prostate cancers develop. *MSMB* as a serum marker and was associated with increased diagnoses of prostate cancer in a case-controls study of ∼1,200 individuals (53). Another case-control study found that serum *MSMB* was not significantly different between cancer and normal groups but was predictive when controlling for age and PSA levels (59). Coupled the results of our analysis, we contend that *MSMB* loss is a marker for metaplasia programs that are one of the earliest markers of de-differentiated epithelial cells, which proceed the morphologic changes necessary for the diagnosis of malignant and pre-malignant lesions. Finally, the observation that DEGAS hazard correlates with total RNA expression is important because recent analysis of the TCGA involving 6590 patients across 15 cancers has shown that total RNA content of a tumor is positively correlated with a poor prognosis (60). Moreover, even though the highest risk region of normal tissue expresses more total RNA, both KLK3 and MSMB expression is lost.

Transcription factors *SNAI1*, *SNAI2*, *ZEB1*, *ZEB2*, *TWIST1*, and *TWIST2* are genes commonly discussed in the epithelial-to-mesenchyme transition (61). Their absence in this analysis was conspicuous. *SNAI2* is the only TF that is upregulated in the high-risk normal tissue of the first dataset (P = 2.60×10^-5^, log2FC = 0.50) and the Erickson et al. BG risk rank 1 (P = 2.30×10^-6^, log2FC = 0.25). This same gene is down-regulated in the Erickson et al BG risk rank 4 (P = 3.1×10^-33^, log2FC = -0.98). Transcription factors, generally, may be lowly expressed genes: They have high affinity for target DNA sequences and operate in the nucleus. They may not need high levels of transcripts to function, and thus, be liable to dropout events. For example, there are 2543 probes in the normal tissue ST array from the first dataset. The total number reads for SNAI1, SNAI2, TWIST2, ZEB1, and Z2B2 across the sample are 42, 2606, 17, 856, 1348, and 980 respectively. This is an average of 0.017, 1.023, 0.0067, 0.3366, 0.53, and 0.385 transcripts detected at each spot. In comparison, the cytoskeletal/structure proteins have much higher expression: Vimentin (VIM), Fibronectin (FN1), β-catenin (CTNNB1), and β-actin (ACTB) have 14753, 9233, 6369, and 79780 reads respectively. The transcription factors are low-expression genes, and direct study of them with high-dropout technologies like ST is likely not appropriate, and instead pursued with more sensitive techniques like fluorescence probe techniques.

### Cell-type decomposition of external dataset and cell-specific differential expression associated with poor progression-free survival

The authors who created this validation dataset were analyzing copy-number variations in the spatial transcriptomics data. They identified that benign tissues have copy number variations that share phylogeny with tumors within the same prostate. Like in the discovery dataset, DEGAS highlighted benign morphology glands that were associated with progression-free survival, and these glands correlated with the number of CNV changes identified by the original authors (SCC = 0.652). Risk scores increased with increasing grade of prostate cancer, aligning with intuition, with GG4 glands being the highest risk and grade. The normal tissue contained four modes of risk: These ranks had distinct spatial distributions. DEGAS does not use any spatial information, so these results are striking and suggest a clonal/evolutionary nature to these groups of cells, aligning with what the original authors found.

Cell-type decomposition at each ST spot revealed the striking observation that single-cell annotations for cancer cells were attributed to regions of benign tissue morphology. This raises an important concern as to how to approach annotating single cell datasets: Rather than annotating spatial data with single cell data, researchers may want to do the reverse as well, transferring histological labels from ST data to SC data. Then, using DEGAS to provide disease association scores can further refine cell clusters and provide clinically-oriented hypotheses. Cell-type specific differential gene expression associated with DEGAS hazard revealed several interesting observations. *MSMB* and *NEFH* were again identified as being negatively associated with progression-free survival. *MSMO1* (*SC4MOL*) is also downregulated in the luminal epithelial cells. Deficiencies in this gene have a causative role in a congenital disorder that involves hyperproliferation and differentiation of keratinocytes by regulation of *EGFR* (62). *AOC1* is inversely associated with risk: This gene has a putative tumor suppressor role in prostate cancer cell lines (63) and seems to have the opposite effect in colon cancer cell lines (64), though these organs come from different embryological germ layers. The basal epithelia can be identified by expression of *S100A11* (42), and C-SIDE analysis revealed that this gene was associated with higher DEGAS risk scores. The basal cells also show downregulation of *CKB*, which has been shown to mediate the EMT and metastasis in prostate cancer mouse xenografts (65). Basal epithelia have shown the capacity to initiate adenocarcinomas where the majority of tumor cells have luminal phenotypes, acting like cancer stem-cells (66). Many genes positively associated with risk in the *ERG* –/+ tumor tissues are ribosome-associated proteins (e.g., *RPL37A*, *RPL23*, *RPS24*).

## Conclusion

With the highly subjective nature of single-cell annotations, tools like DEGAS are essential to add nuance to clustering methods, especially when it is challenging to further separate large clusters of cells (e.g., tumor and epithelial cells) that exhibit a continuum of changes. In our experiments, we demonstrate that identifying significant differences in expression is difficult without a clinically-oriented hypothesis generation tool to pinpoint disease-associated cells. Furthermore, we highlight the challenges of building single-cell annotation atlases by integrating single-cell and transcriptomics data, as these atlases often disregard cell morphology and spatial arrangements. The ERG-negative tumor cells attributed to high-risk benign glands in our analysis may represent distinct cell states, rather than being genuine ERG-negative tumor cells.

We show that MSMB loss in luminal epithelial cells in morphologically benign epithelial cells is associated with poor progression-free survival. Previous studies have linked MSMB loss in prostate cancer to recurrence. Our observed associations between MSMB loss and increased cancer RNA signatures in normal epithelia contribute to the current understanding of MSMB. We hypothesize that this gene is lost during metaplastic changes preceding morphological alterations. Future research will focus on validating a gene panel for immunohistochemistry (IHC) in benign prostate biopsy specimens to assess the efficacy of predicting prostate cancer incidence. This is crucial, as current preventative measures are imprecise and often lead to overtreatment, and it could provide researchers with a tool to study early carcinogenesis. Utilizing DEGAS to examine normal biopsies may offer similar insights into other cancers, and we have showcased this tool’s utility for rapid, clinically-oriented hypothesis generation.

---

**Supplemental Figure 1:**
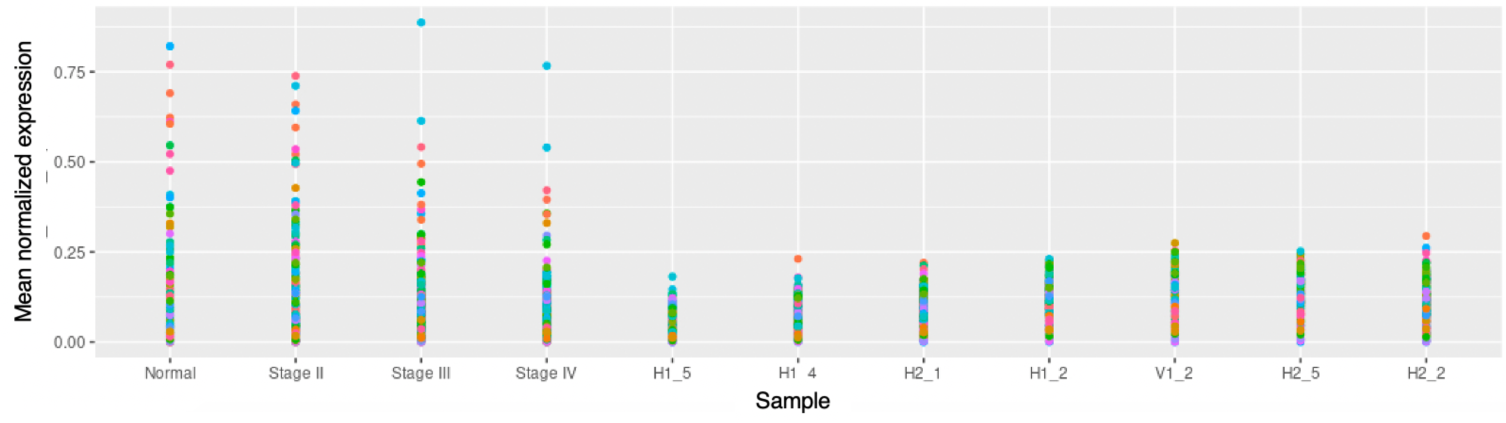
Independent feature selection DEGAS of datasets is necessary. To analyze the the publicly-available 10x prostate samples (Normal, Stage III, III, and IV), the top 200 most variable genes in the TCGA were used as features in the DEGAS model. These same 200 genes have different distributions, and some do not appear in the Erickson data, demonstrating potential differences in biology and/or sample preparation.

**Supplemental Figure 2:**
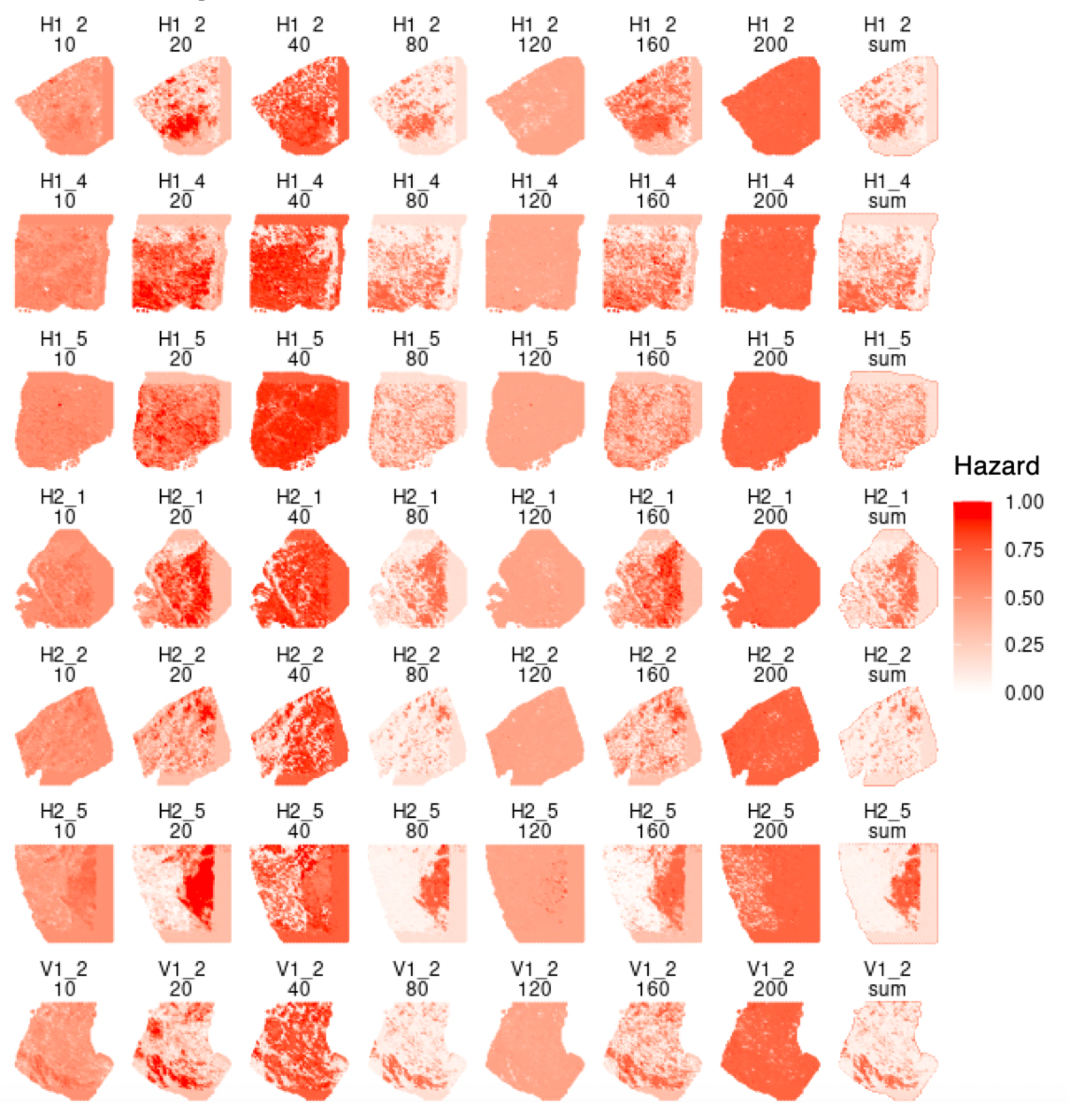
Effect of feature selection on hazard output. Similar patterns emerge across different levels of feature selection. Taking the sum of predicted hazards across increasing features sets provides a regularized DEGAS output.

**Supplemental Figure 3:**
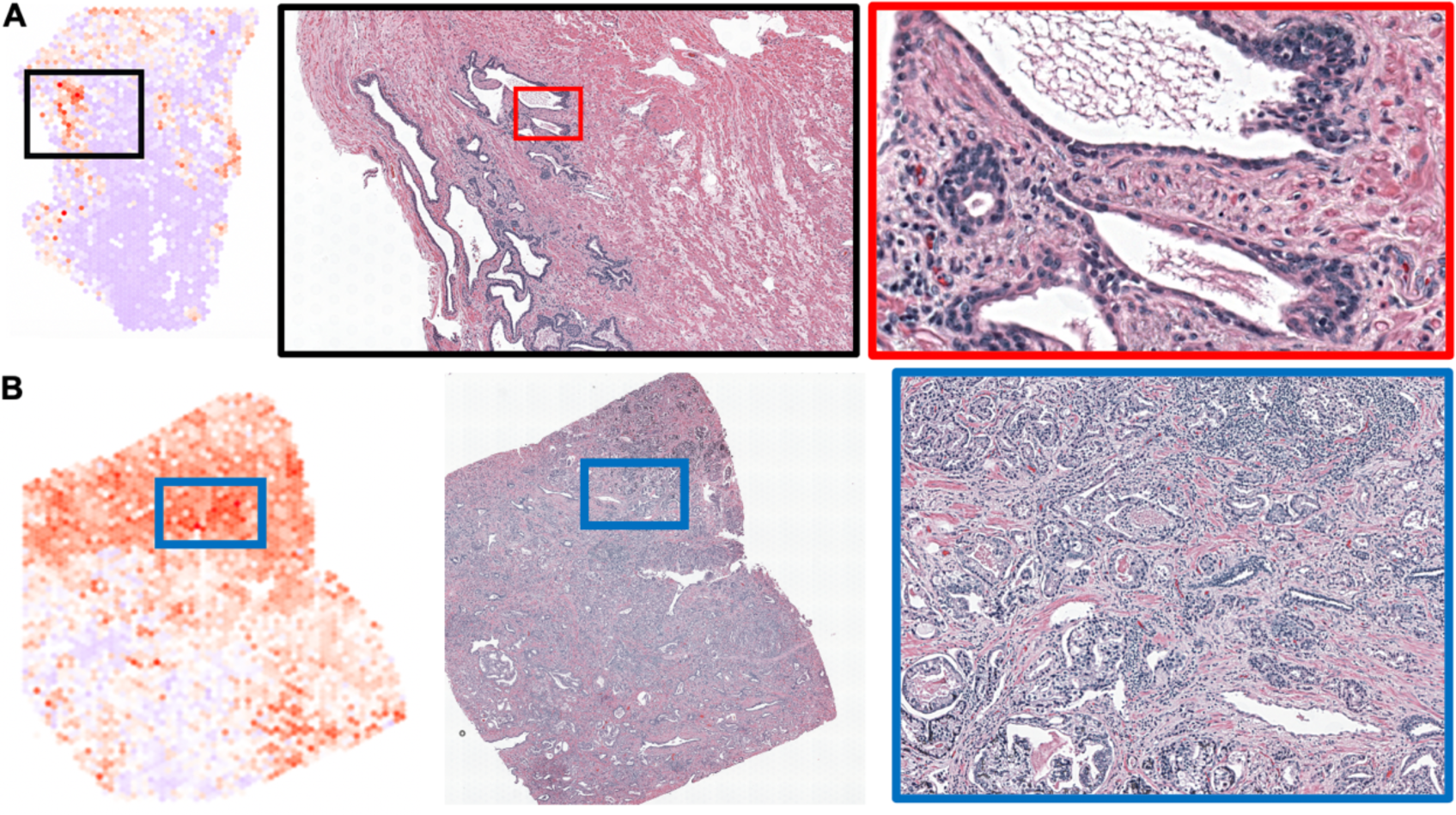
Normal tissue and Stage IV adenocarcinoma high risk regions. Regions of high-risk morphologies visualized. **A.** Highest risk glandular region in the normal prostate tissue. There are no obvious signs of atypia. **B.** Highest risk regions tend to have back-to-back glands or completely dissociated single cancer cells, indicating higher Gleason scores.

**Supplemental Figure 4:**
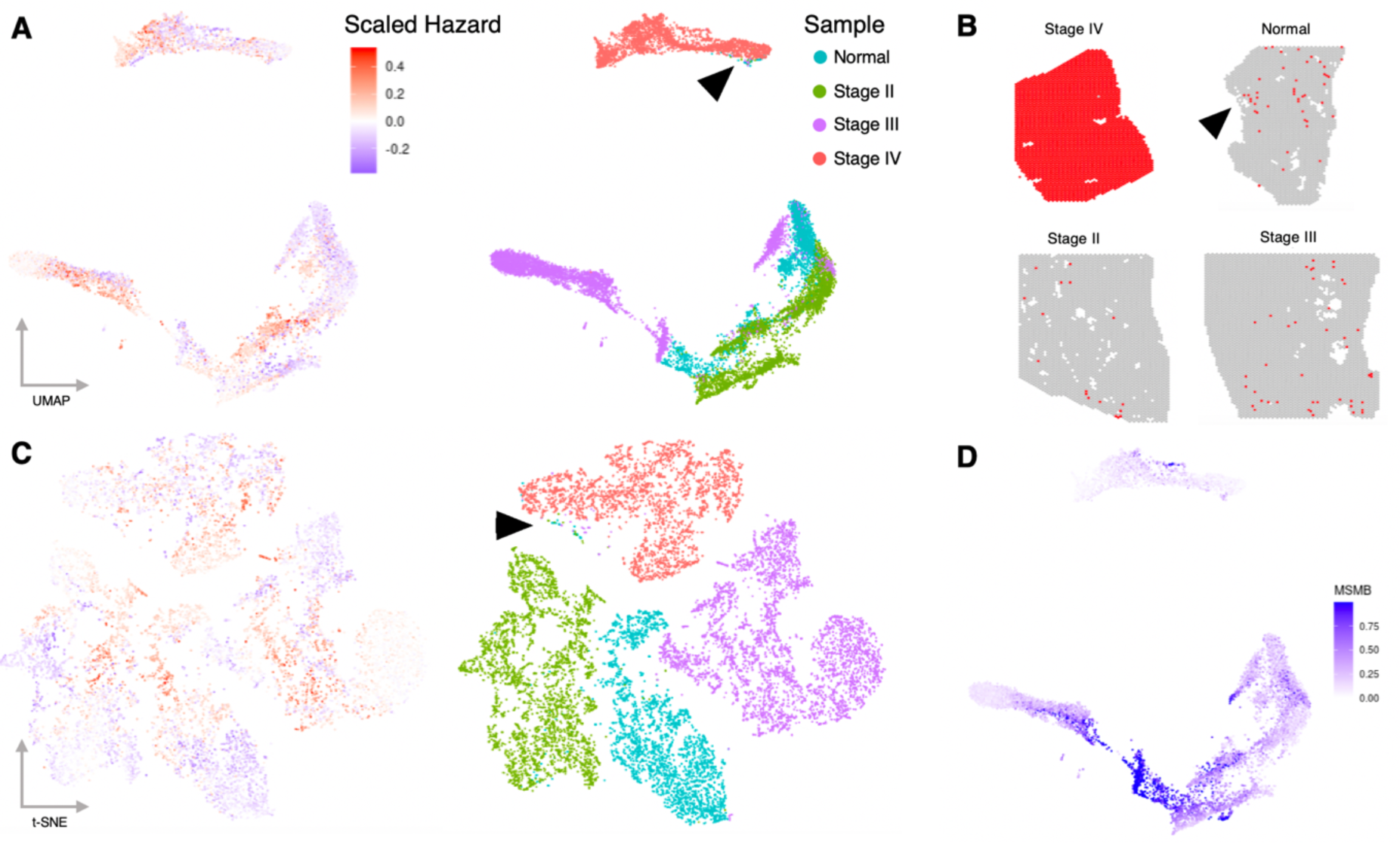
UMAP and t-SNE of four prostate samples. UMAP **(A)** and t-SNE **(C)** of top 3000 most variable genes shows in the four prostate ST samples. Black arrows in **A** and **C** highlight regions of high dimensional similarities between the normal, Stages II, III, and IV cancers. Red arrow shows the high-risk glandular region as being. **B.** Clusters indicated by black arrow in A, which is mostly Stage IV ST spots, plotted in the original ST data. All spots from Stage IV sample appear in the cluster. Spots from the other samples mostly coincide with the high-risk regions identified by DEGAS (black arrow next to normal sample indicates general area of highest-risk glands). High dimensional clusters do not represent the full continuum of DEGAS risk outputs. **D.** MSMB expression plotting in UMAP shows low expression tends to align with higher-risk regions **(A)**.

**Supplemental Figure 5:**
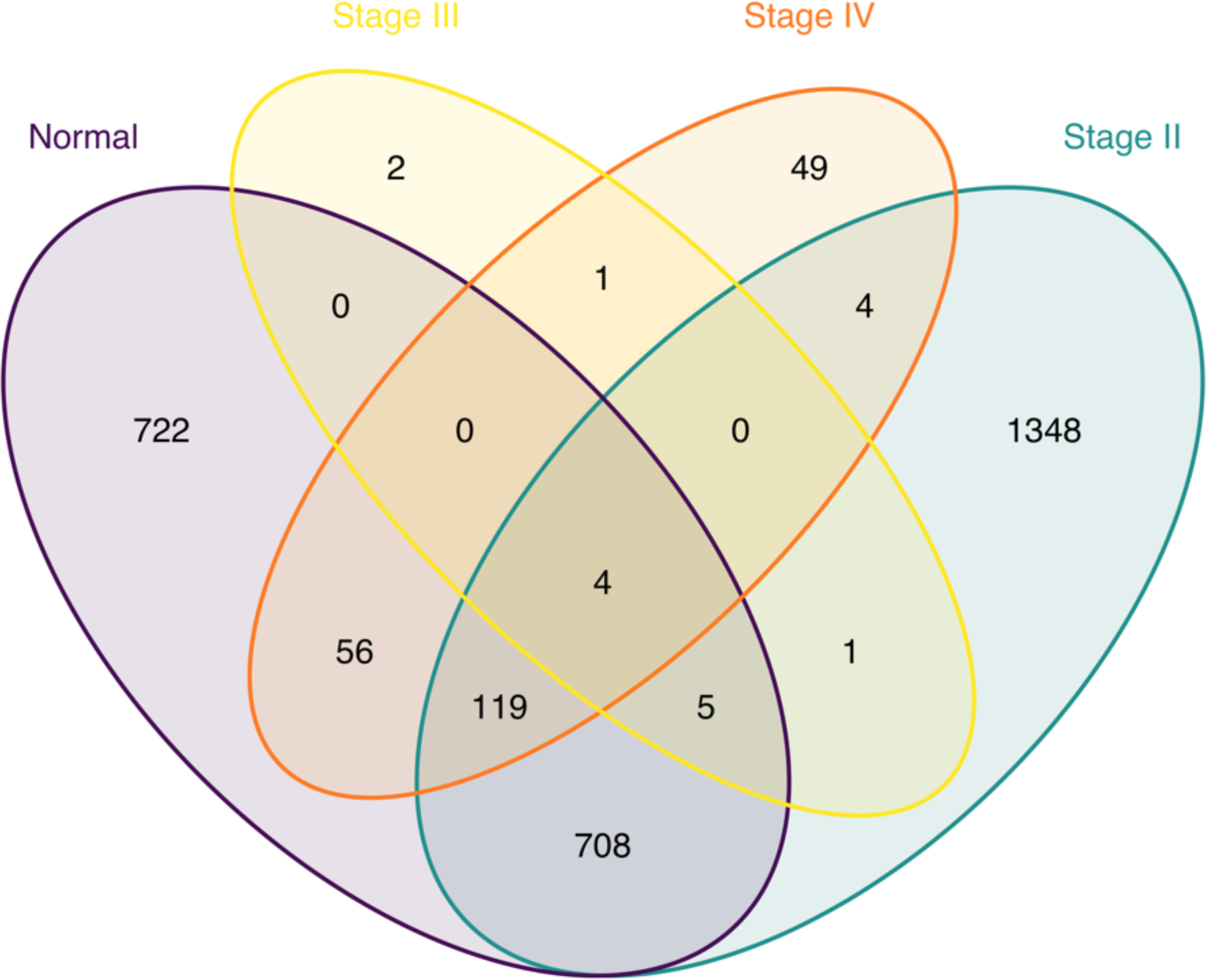
Overlap in upregulated genes in high hazard regions across all four prostate ST samples. Wilcoxon test was used to generate p-values for differential expression of high and low hazard ST spots for each sample. Among the genes with B-H FDR-adjusted p-values less than 1E-2, up to 100 genes were input to Toppgene for functional enrichment. The overlap between GO ontology categories of *Molecular Function, Biological Process, Cellular Component, Human Phenotype, Mouse Phenotype, Pathway, Interaction,* and *Disease* are visualized with this Venn diagram.

**Supplemental Figure 6:**
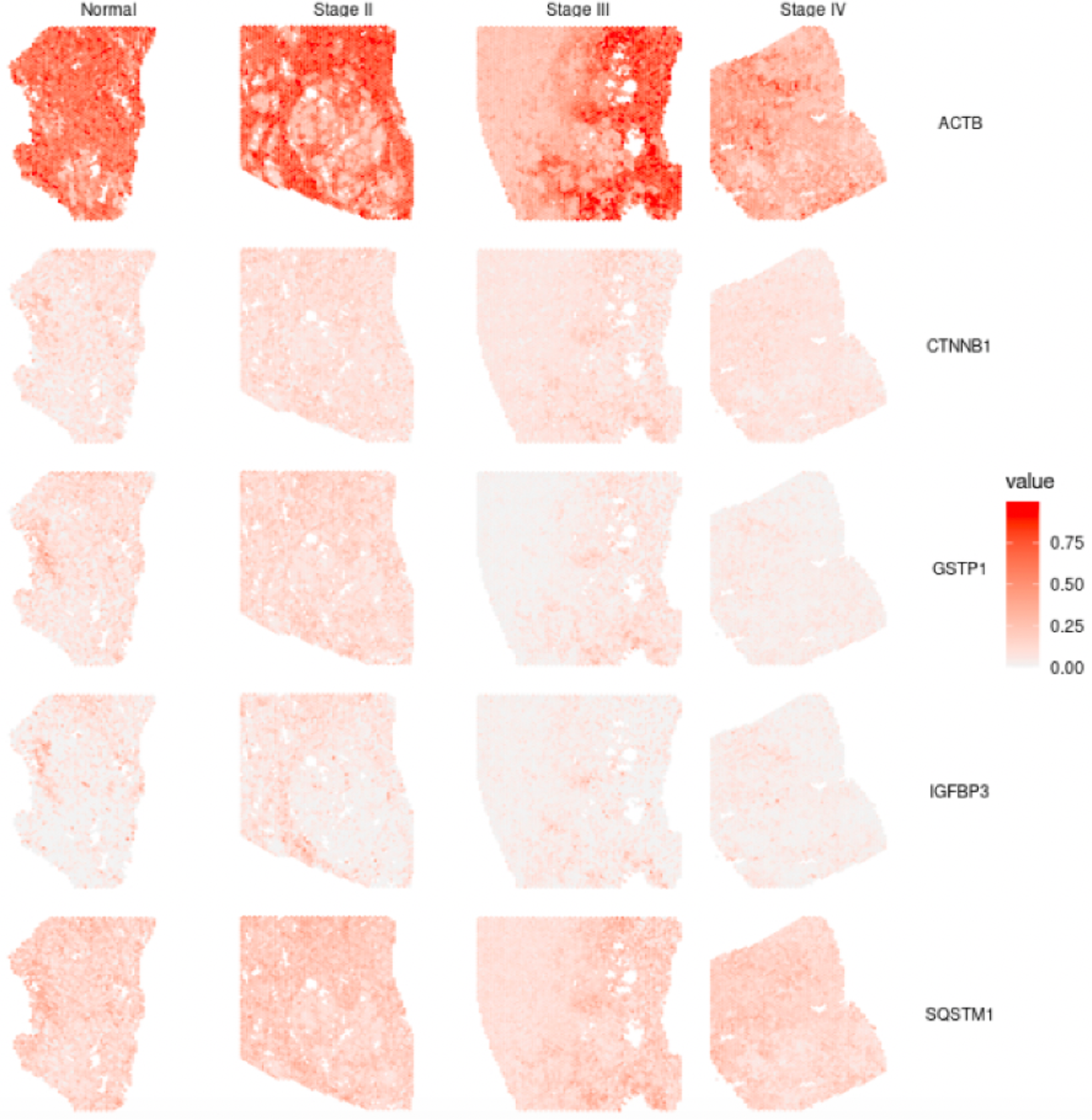
High Grade PIN genes from normal glands. Genes contributing to enrichment for High Grade Prostatic Intraepithelial Neoplasia from the high-risk glandular tissue from the normal sample, plotted on all prostate samples. Beta-catenin 1 expression is low in the low-risk glands. Expression is higher in the stroma and in the highest risk glandular areas.

**Supplemental Figure 7:**
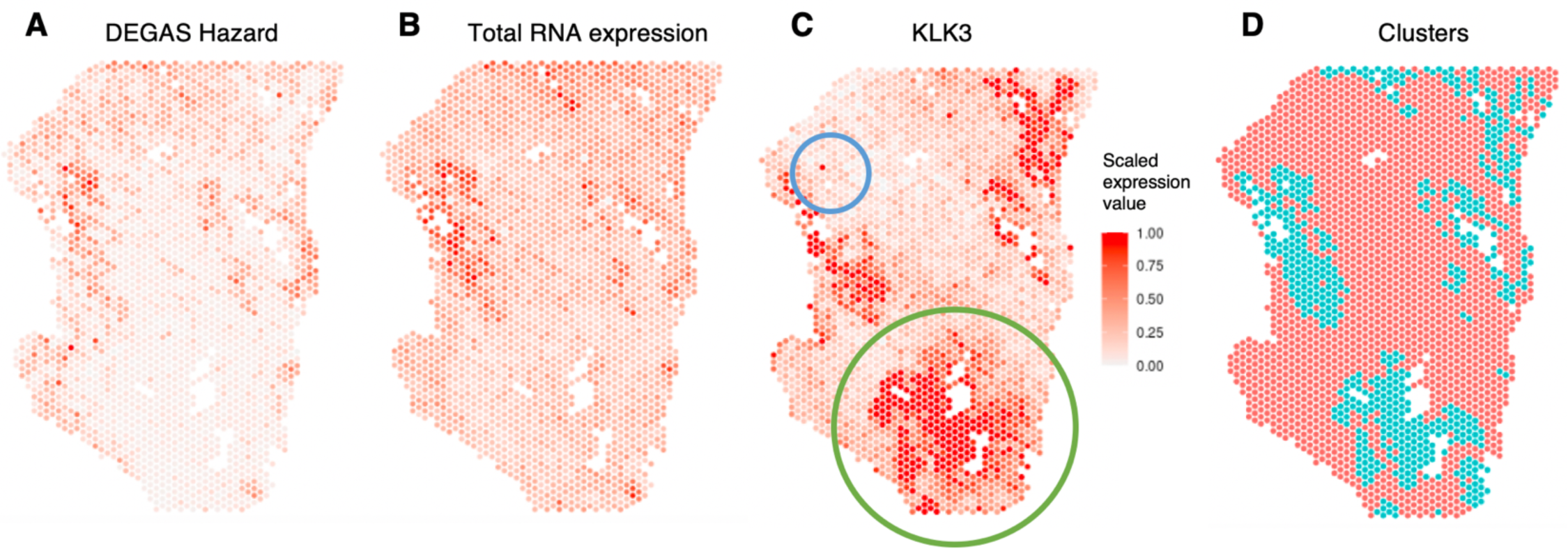
DEGAS hazard output correlates with total RNA expression, the low-risk glandular regions express more KLK3. **A.** DEGAS-identified high-risk regions tend to align with higher levels of (**B**) total RNA expression. The Spearman correlation between these variables is 0.546. **C**. KLK3 is the protein measured as “Prostate serum-antigen” or “PSA” and is expressed by prostatic glandular tissues (including cancerous tissue). Some of the highest-risk regions show no *KLK3* expression (blue circle), and the lowest-risk regions show high *KLK3* expression (green circle), despite being glandular tissue. **D**. K-means cluster (K=2) show the regions of glandular (blue) and stromal prostate tissues (red).

**Supplemental Figure 8:**
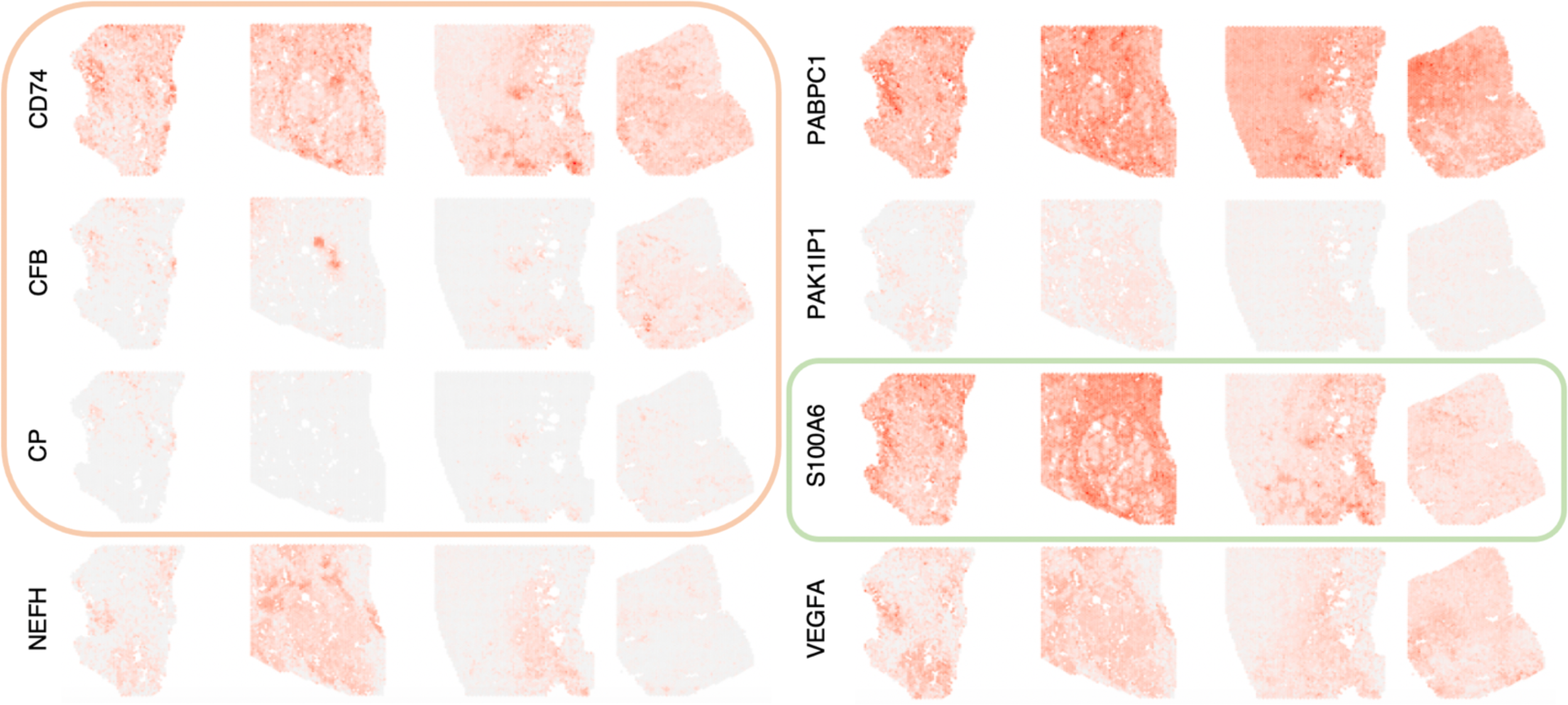
Highlighted genes from proteomic integration visualized on ST. Visualization of RNA expression across all four prostate samples for genes. Genes in the orange box are not detected in normal proteomic data, but more highly expressed in cancer proteomics. *S100A6*, green box, is moderately expressed in normal but not cancer tissue. *VEGFA* is downregulated in the high-risk normal glands, and consistently negatively correlated with DEGAS risk scores. *PAK1P1* is also downregulated, but the correlation with DEGAS risk scores is inconsistent.

**Supplemental Figure 9:**
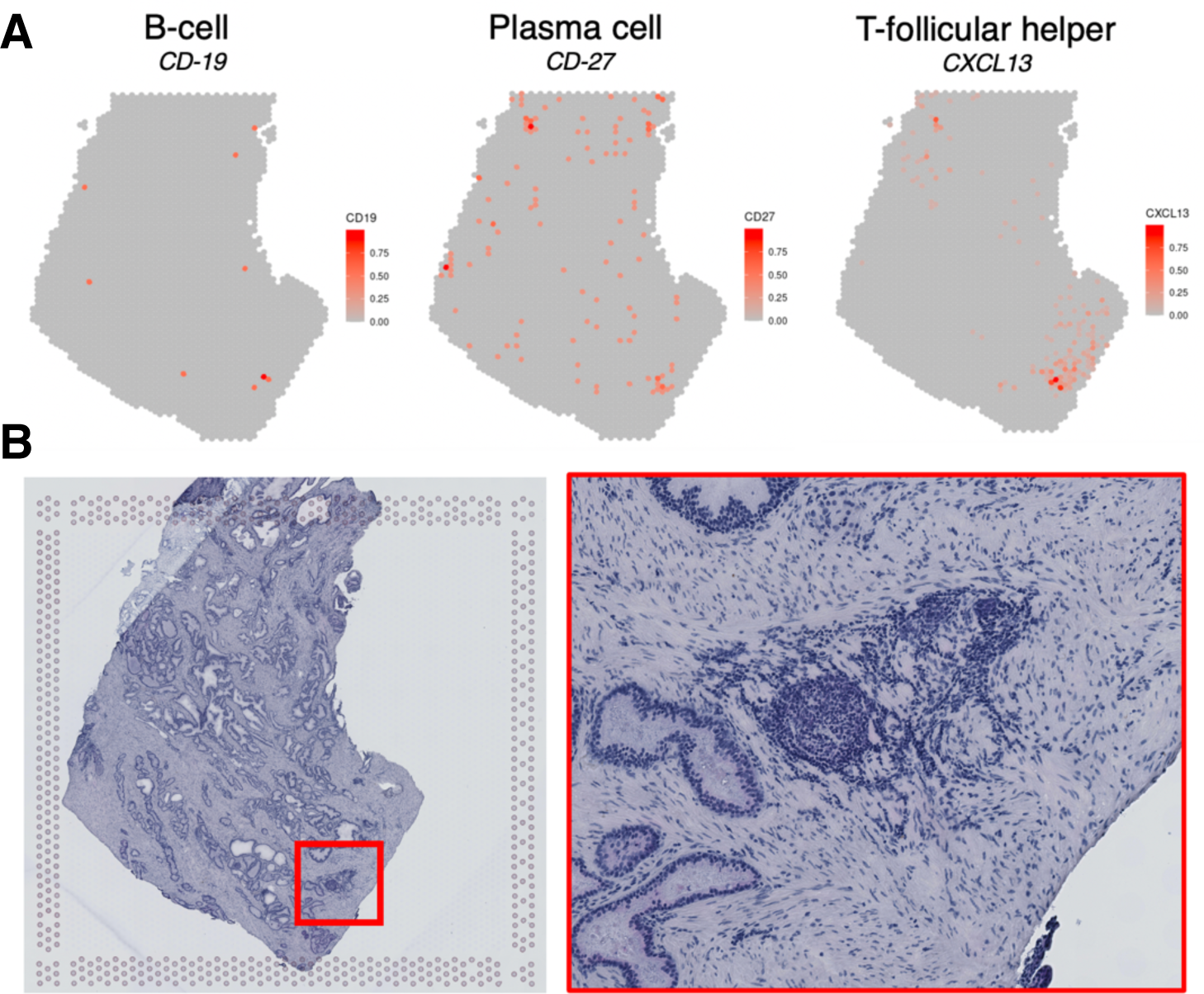
B-cell group from RCTD analysis aligns with histology of inflamed epithelial structure. **A.** B-cell enrichment from RCTD aligns with markers for B cells (CD-19), Plasma cells (CD-27), and T-follicular helper cells (CXCL13). **B.** The corresponding location in the histopathology shows an inflamed epithelial structure.

